# Inducible transgene expression in PDX models *in vivo* identifies KLF4 as therapeutic target for B-ALL

**DOI:** 10.1101/737726

**Authors:** Wen-Hsin Liu, Paulina Mrozek-Gorska, Tobias Herold, Larissa Schwarzkopf, Dagmar Pich, Kerstin Völse, Anna-Katharina Wirth, M. Camila Melo-Narváez, Michela Carlet, Wolfgang Hammerschmidt, Irmela Jeremias

## Abstract

Clinic-close methods are not available that prioritize and validate potential therapeutic targets in individual tumors from the vast bulk of descriptive expression data. We developed a novel technique to express transgenes in established patient-derived xenograft (PDX) models *in vivo* to fill this gap. With this technique at hand, we analyzed the role of transcription factor Krüppel-like factor 4 (KLF4) in B-cell acute lymphoblastic leukemia (B-ALL) PDX models at different disease stages. In competitive pre-clinical *in vivo* trials, we found that re-expression of wild type KLF4 reduced leukemia load in PDX models of B-ALL, with strongest effects after conventional chemotherapy at minimal residual disease (MRD). A non-functional KLF4 mutant had no effect in this model. Re-expressing KLF4 sensitized tumor cells in the PDX model towards systemic chemotherapy *in vivo*. Of major translational relevance, Azacitidine upregulated KLF4 levels in the PDX model and a KLF4 knockout reduced Azacitidine-induced cell death, suggesting that Azacitidine can regulate KLF4 re-expression. These results support applying Azacitidine in patients with B-ALL to regulated KLF4 as a therapeutic option. Taken together, our novel technique allows studying the function of dysregulated genes in a highly clinic-related, translational context and testing clinically applicable drugs in a relevant pre-clinical model.

## Introduction

Tumor cells are characterized by multiple alterations at the level of mRNA and protein expression. While non-genomic alterations are easily identified by techniques such as RNA sequencing or proteomics (1–3), functional consequences of these alterations are generally uncertain and only a small minority might play an essential function for growth and maintenance of the tumor. Identifying essential alterations is of major clinical importance as they represent putative targets for treatment (1–4). Clearly, techniques are missing to validate alterations for their functional consequence, but we established a clinic-close method to identify alterations with life-sustaining function in individual tumors *in vivo*. As a model system, we chose patient-derived xenografts (PDX), which overcome many of the limitations associated with the use of conventional cell line models or primary patient tumor cells (5–7). PDX faithfully recapitulate genetic and phenotypic characteristics of their parental tumors and their pre-clinical value for drug-testing and biomarker identification is well established (8, 9). Thus, orthotopic PDX models represent the most clinical-related approach up to date for studying individual tumors (6, 10). We studied B-cell precursor ALL (B-ALL), the most frequent malignancy in children, which requires better treatment options, especially upon disease relapse (11). To mimic the situation of patients, established tumors should already exist in the preclinical surrogate animals at the moment of molecular manipulation, which requires inducible *in vivo* expression systems. While these techniques are well established in most other tumor models, inducible orthotopic PDX models have not been reported to our knowledge. We generated an inducible expression system in PDX models in mice *in vivo*. We applied and tested our novel method using an exemplary signaling molecule, the transcription factor KLF4, which is implicated in stress-responsive regulation of cell cycle progression, apoptosis and differentiation as well as stemness and pluripotency (12–18). KLF4 is downregulated in numerous cancers, but upregulated in others, and abnormal KLF4 expression might reflect oncogenic or tumor suppressor functions depending on the cellular context, tumor type, sub-type and stage (13, 16, 19–26). KLF4 is frequently deregulated in T-ALL and B-cell tumors (27–33), and reportedly downregulated in pediatric B-ALL (34–37). While KLF4 downregulation has been implicated in B-ALL leukemogenesis (38), its functional role in established patient B-ALL cells *in vivo* is uncertain. We developed a tetracycline-inducible expression system to re-express KLF4 in PDX B-ALL cells *in vivo*. We demonstrate here that KLF4 expression reduces tumor growth and increases chemotherapy response in this tumor model. With the aid of a CRISPR/Cas9-mediated KLF4 knockout in PDX cells we further demonstrate that Azacitidine exerts its anti-tumor effect by upregulating KLF4, supporting our interpretation. Our data demonstrate that inducible gene expression in PDX models is feasible, can characterize the contribution of selected genes to tumor maintenance and delivers valuable information regarding therapy response. Our results reveal that KLF4 is an interesting therapeutic target in B-ALL, supporting the use of KLF4 regulating drugs in clinical trials of B-ALL.

## Results

### KLF4 blocks EBV-induced transformation of B-cells *in vitro*

We focused on studying KLF4 as a prime example, because we found that KLF4 is severely downregulated in primary pediatric B-ALL cells after therapy (39) (*Supplementary Figure 1A*), an observation that complements published data on low KLF4 expression in pediatric B-ALL (34–37). In adult ALL samples, the aggressive Ph+-like B-ALL sub-group showed similarly low expression levels of KLF4 as T-ALL (23) (*Supplementary Figure 1B*). In line with ALL, strong and persistent downregulation of KLF4 mRNA levels was detected during EBV-induced B-cell transformation (40) (*Supplementary Figure 2A*), an *in vitro* model of B cell oncogenesis.

We performed reverse genetics to decipher the role of KLF4 in certain B-cell tumors. To cover both tumor formation and tumor maintenance, we started with the *in vitro* surrogate model of EBV-induced B-cell transformation that reprograms B-lymphocytes into B-blasts (Figure 1A) (41–43). Two different variants of KLF4, namely wild type KLF4 (wtKLF4) and a variant form of KLF4 (mutKLF4) were cloned into recombinant viruses. Upon infection, they co-express wtKLF4 or mutKLF4 together with viral genes in primary B-lymphocytes (*Supplementary Figure 2B*). mutKLF4 lacks the two C-terminal zinc fingers representing the DNA binding domain, which disables KLF4’s basic functions as transcription factor, but protein-protein interactions of KLF4 are not affected (*Supplementary Figure 2B*).

**Fig. 1.**
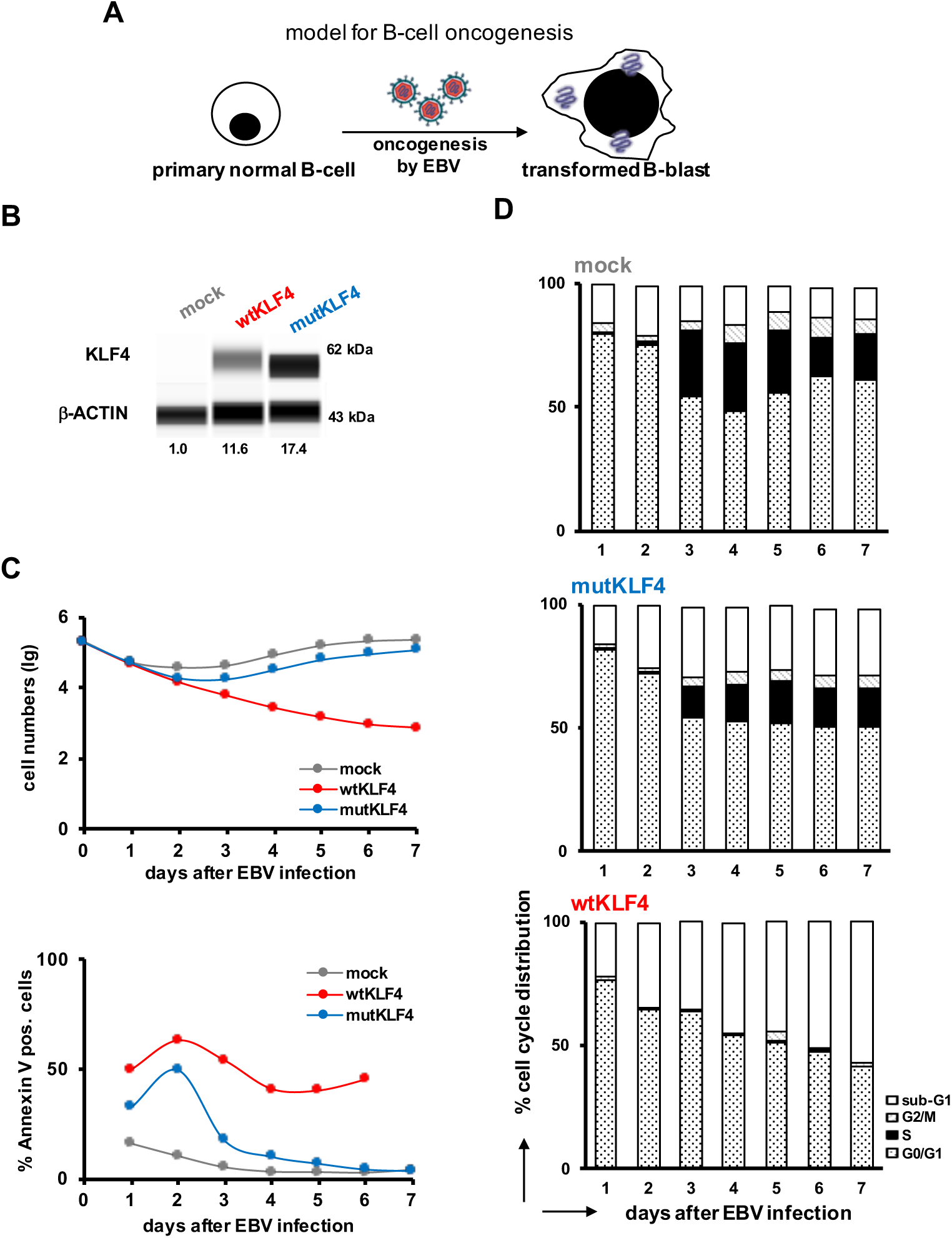
Ectopic expression of KLF4 blocks EBV-mediated oncogenesis. 2×10^5^ normal naïve primary B-cells were infected with EBV *in vitro* to induce their transformation. Either the wildtype EBV virus was used which contained all viral genes required for oncogenesis (mock); alternatively, EBV additionally expressed wild type KLF4 in normal B-cells (wtKLF4) or a mutant KLF4 derivative, lacking the DNA-binding domain (mutKLF4). After EBV infection, B-cells were harvested every day for a total of 7 days. **A** Experimental design **B** Capillary immunoassay of KLF4, two days after infection with the indicated EBV viruses. *β*-actin was used as loading control. **C** Cell numbers were quantified by flow cytometry (upper panel) and apoptosis measured using Annexin V staining (lower panel). **D** Cell cycle analysis; daily, an aliquot of cells was incubated with 5-Bromo-2’-deoxyuridine (BrdU) for 1 hour prior to harvest. The cells were analyzed by flow cytometry after permeabilization and staining with a BrdU specific antibody. The percentage of cells in the different phases of the cell cycle are indicated. One representative experiment out of two is shown using sorted naïve normal B-cells from two individual donors.

Naïve human B-lymphocytes were infected with EBV to induce transformation into B-blasts by viral oncogenes (Figure 1A, mock). To study the influence of KLF4 for the process of transformation, EBV was genetically modified and used as a vector shuttle to express wtKLF4 or mutKLF4 together with the viral oncogenes in resting B-lymphocytes that undergo EBV-induced oncogenesis (Figure 1B, *Supplementary Figure 1C*). While mutKLF4 expression showed no major effects compared with mock EBV-infected B-cells, persistent wtKLF4 levels completely abrogated EBV-induced B-cell transformation, disabled EBV-induced B-cell proliferation and induced apoptosis in the infected cells (Figure 1C). Cell cycle analysis indicated that KLF4 expression prevented S-phase entry of EBV-infected B-lymphocytes, required for generation and expansion of B-blasts. Thus, KLF4 acts as a cell cycle inhibitor in the model of *in vitro* EBV-induced oncogenesis of normal B-cells into B-blasts.

In conclusion, persistent expression of KLF4 blocks human B-cell transformation in the model of EBV-induced oncogenesis of normal B-cells into B-blasts. This finding is in line with published data that suggest a role of KLF4 as tumor suppressor in BCR-ABL-mediated transformation of murine pre-B-cells(38). In both models of B-cell malignancies, murine leukemia and human lymphoma, ectopic expression of KLF4 was incompatible with oncogenesis.

### Inducible transgene expression in PDX cells *in vivo*

We asked whether KLF4 is functionally relevant for established B-cell tumors *in vivo* and whether it might represent a putative therapeutic target. As a model of clinic-close functional genomic studies in established tumors, we used orthotopic PDX from two children with relapsed B-cell precursor ALL (clinical data for both patients are published in Ebinger et al. (39)). Similarly to pediatric (34–37) and adult (*Supplementary Figure 1B*) primary B-ALL samples, both PDX B-ALL models revealed low KLF4 mRNA and protein levels (Figure 2A and B).

**Fig. 2.**
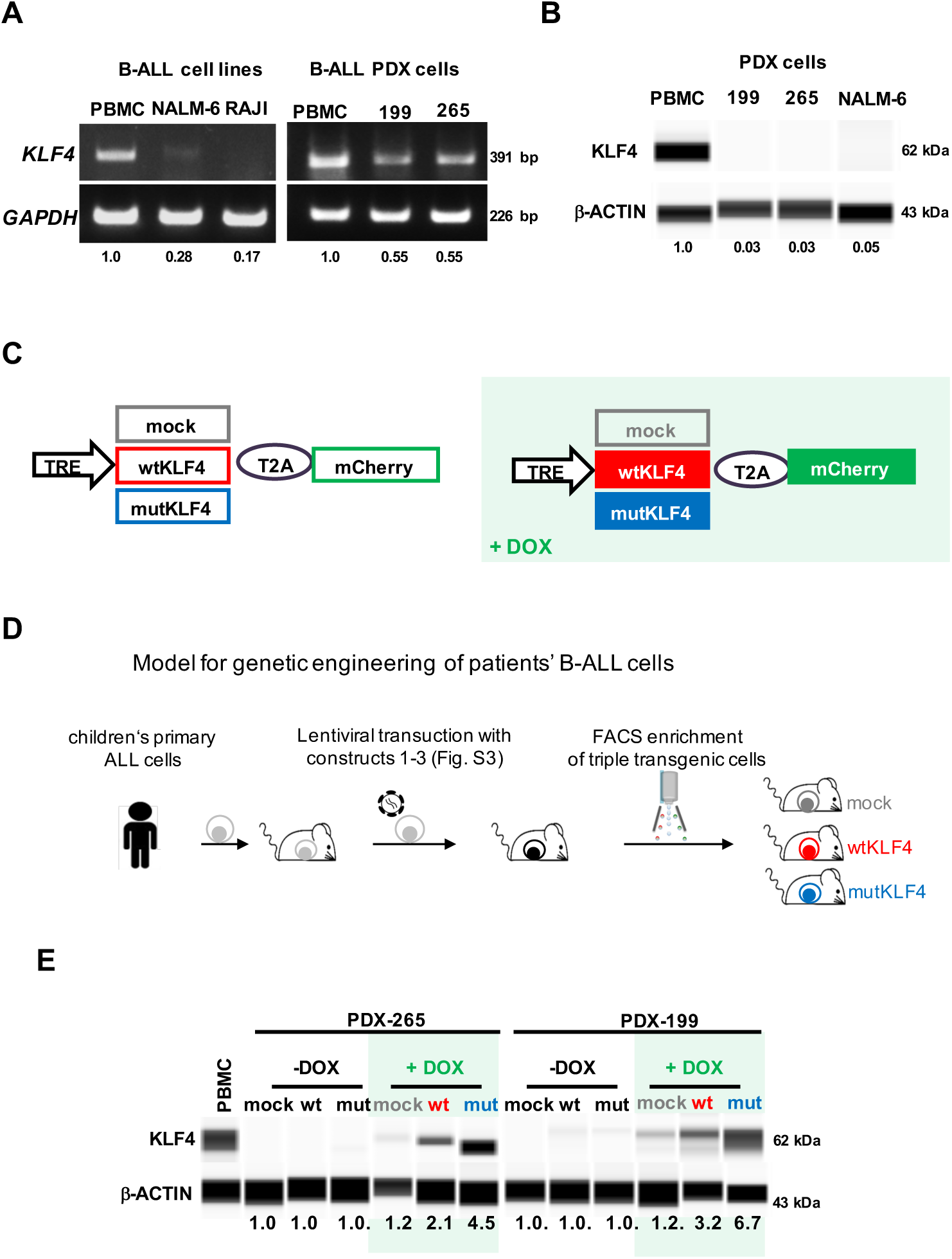
Re-expression of KLF4 in PDX ALL cells using a tet-on inducible system. **A**,**B** KLF4 is downregulated in B-ALL cell lines, B-cell lymphoma cell lines and B-ALL PDX cells as compared to peripheral blood mononuclear cells (PBMC). (**A**) *KLF4* mRNA was determined by RT-PCR using GAPDH as loading control; (**B**) KLF4 protein levels were analyzed by capillary immunoassay using *β*-actin as loading control. **C** KLF4 expression vector. The TRE promoter drives expression of KLF4, either as wildtype (wt)KLF4 or a mutated (mut)KLF4 sequence devoid of two C-terminal zinc finger motifs comprising the DNA-binding domain (see *Supplementary Figure 2B*), each linked by a T2A peptide to the fluorochrome mCherry as molecular marker; mock (empty vector only encoding the mCherry fluorochrome) was used as a control. Addition of Doxycycline (light green background throughout all Figures) leads to expression of mock (grey), wtKLF4 (red) or mutKLF4 (blue; colors identical throughout all Figures) and mCherry (green). **D** Experimental design: Primary B-ALL cells from patients were transplanted into immunocompromised mice to generate PDX models; PDX cells were consecutively transduced with three lentiviral constructs to express rtTA3 together with luciferase, tetR and either mock, wtKLF4 or mutKLF4 (vectors are detailed in panel C and *Supplementary Figure 3A*). Following passaging through mice for amplification, transgenic PDX cells were enriched by flow cytometry based on constitutively expressed fluorochromes (mTaqBFP for the rtTA3-luciferase construct, iRFP720 or T-Sapphire for the tetR construct and Venus for the KLF4 constructs). Triple-transgenic cells were used for all further in vivo experiments. **E** PDX ALL-265 and PDX ALL-199 infected as indicated in (**D**) were cultured *in vitro* with or without addition of DOX for 48h and KLF4 protein expression was analyzed by capillary immunoassay using peripheral blood mononuclear cells (PBMC) from healthy donors and *β*-actin as controls. Representative analysis from triplicates are shown.

We developed a novel method to conditionally express KLF4 in PDX B-ALL cells. Towards this end we engineered a tetracycline-response system to study characteristics of PDX models independently of tumor cell transplantation and at different time points, e.g., before, during and after treatment and at minimal residual disease. PDX cells were consecutively transduced with three lentiviral constructs (Figure 2C and D and *Supplementary Figure 3A*) to express (i) the tet activator (rtTA3) together with luciferase to allow *in vivo* bioluminescence imaging; (ii) the tet repressor (tetR) together with one of two different fluorochromes as molecular markers for competitive *in vivo* assays; and (iii) a mock, wtKLF4 or mutKLF4 gene (*Supplementary Figure 2B*) in conjunction with the fluorochrome mCherry and under the control of the tet-responsive element (TRE)(44) (Figure 2D and *Supplementary Figure 3A*). We chose a low transduction rate of 1% for each vector to integrate only single transgene copies per cellular genome. Although gene silencing of the CMV promoter is common in leukemic cells but the TRE-regulated mini-CMV promoter did not become epigenetically repressed in our PDX models (data not shown). Doxycycline (DOX) treatment (indicated by green background in all Figures) of transduced B-ALL cells induced expression of KLF4 together with mCherry, which serves as a proxy to monitor the co-expression of the wtKLF4 or mutKLF4 proteins (Figure 2E and *Supplementary Figure 3B* and C). Upon DOX withdrawal expression of mCherry together with KLF4 turned back to baseline, demonstrating the reversibility of the system (*Supplementary Figure 3C*). Of note, KLF4 protein expression in the induced state was moderate and did not exceed levels observed in PBMC, avoiding undesired overexpression (Figure 2E). Importantly, feeding mice with Doxycycline (DOX) induced expression of mCherry/KLF4 in PDX B-ALL cells *in vivo* (*Supplementary Figure 3B*), enabling the analysis of KLF4-mediated effects at any given time point in mice. The combination of different marker fluorochromes opened the opportunity to investigate PDX cells with conditional wtKLF4 and control (mock) alleles in the same animal by pairwise competition experiments abrogating animal-to-animal variability.

Taken together, the newly established inducible system of transgene expression in PDX acute leukemias appears operational and represents an attractive method to identify and study individual leukemia vulnerabilities and to test established and future drugs for their anti-leukemic potency *in vivo*.

### Re-expressing wtKLF4 reduces growth and homing of B-ALL PDX cells *in vivo*

In a first step, we determined whether low KLF4 expression levels were essential and required for survival and proliferation of B-ALL tumors *in vivo*. Two PDX B-ALL samples, ALL-265 (Figures 3–5) and ALL-199 (*Supplementary Figure 4*) cells transduced with either mock or wtKLF4 alleles were transplanted into groups of mice. When homing and early engraftment were completed, mice were fed with DOX to induce transgene expression during the exponential growth phase of pre-established PDX B-ALL tumors (Figure 3A and *Supplementary Figure 4A*). Tumor growth of wtKLF4-expressing B-ALL PDX cells was reduced compared with controls as determined by *in vivo* imaging (Figure 3B and *Supplementary Figure 4B*). Consequently, KLF4 re-expression resulted in reduced splenomegaly compared with spleens from mock-bearing mice (Figure 3C and D and *Supplementary Figure 4CD*). KLF4 significantly reduced the fraction of cells in S-phase (Figure 3E and *Supplementary Figure 4E*) and induced cleavage of PARP and Caspase-3 (Figure 3F and *Supplementary Figure 4F*) indicating that KLF4 impaired tumor growth by inducing cell cycle arrest and apoptosis. Comparable results were obtained with B-ALL cell lines (*Supplementary Figure 5*) supporting our observations *in vivo*.

**Fig. 3.**
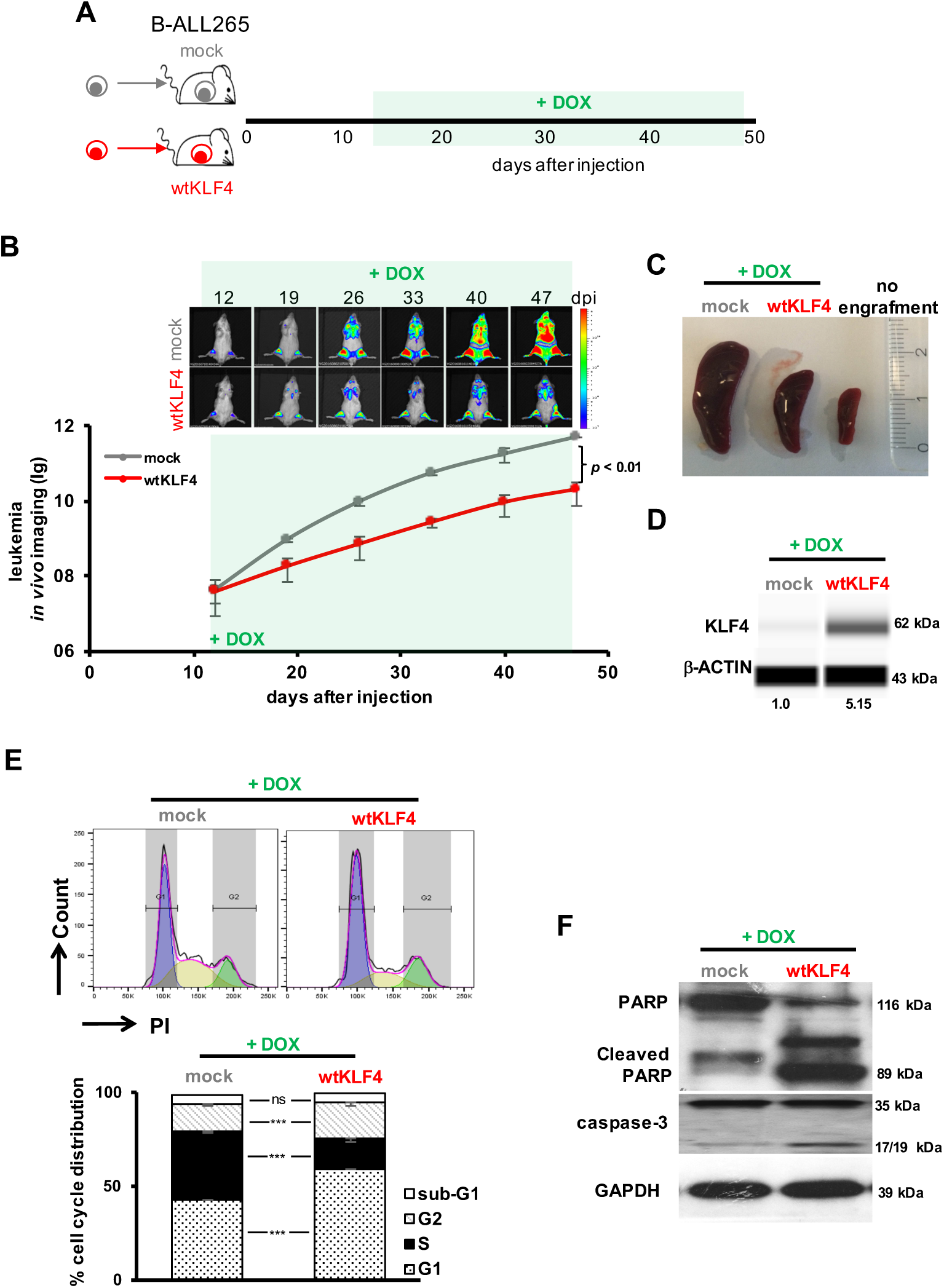
Re-expressing KLF4 inhibits tumor growth through cell cycle arrest and apoptosis in B-ALL PDX cells *in vivo*. **A** Experimental design: 60,000 triple-transgenic, mock or wtKLF4 PDX ALL-265 cells were injected into 6 NSG mice each. After homing was completed and tumors were established, DOX (1mg/ml) was added to the drinking water on day 12 to induce KLF4 expression. On day 47 after cell injection, mice were sacrificed, spleens harvested and the DOX-induced, mCherry positive mock or wtKLF4 expressing population enriched by flow cytometry for further analysis. **B** Bioluminescence *in vivo* imaging displayed as representative images (upper panel) and after quantification of all 6 mice, depicted as mean ± SEM. ** p<0.01 by two-tailed unpaired t test. **C** Representative spleens of mice, using a healthy mouse without leukemic engraftment for comparison. **D** KLF4 protein level of mCherry positive splenic cells was analyzed by capillary immunoassay; *β*-actin served as loading control. Representative analysis from one out of three mice are shown. **E** Cell cycle analysis; 10^6^ mCherry positive cells were stained with Propidium Iodide (PI) and cell cycle distribution was measured by flow cytometry. Upper panel: Representative histograms of mock (n=3) or wtKLF4-expressing (n=3) cells; lower panel: Quantification as mean ± SD is shown. *** p<0.005 by two-tailed unpaired t test; ns = not significant. **F** PARP and caspase-3 cleavage in mock- (n=3) or wtKLF4-transduced (n=3) PDX ALL-265 cells as determined by Western blot; GAPDH was used as loading control.

**Fig. 4.**
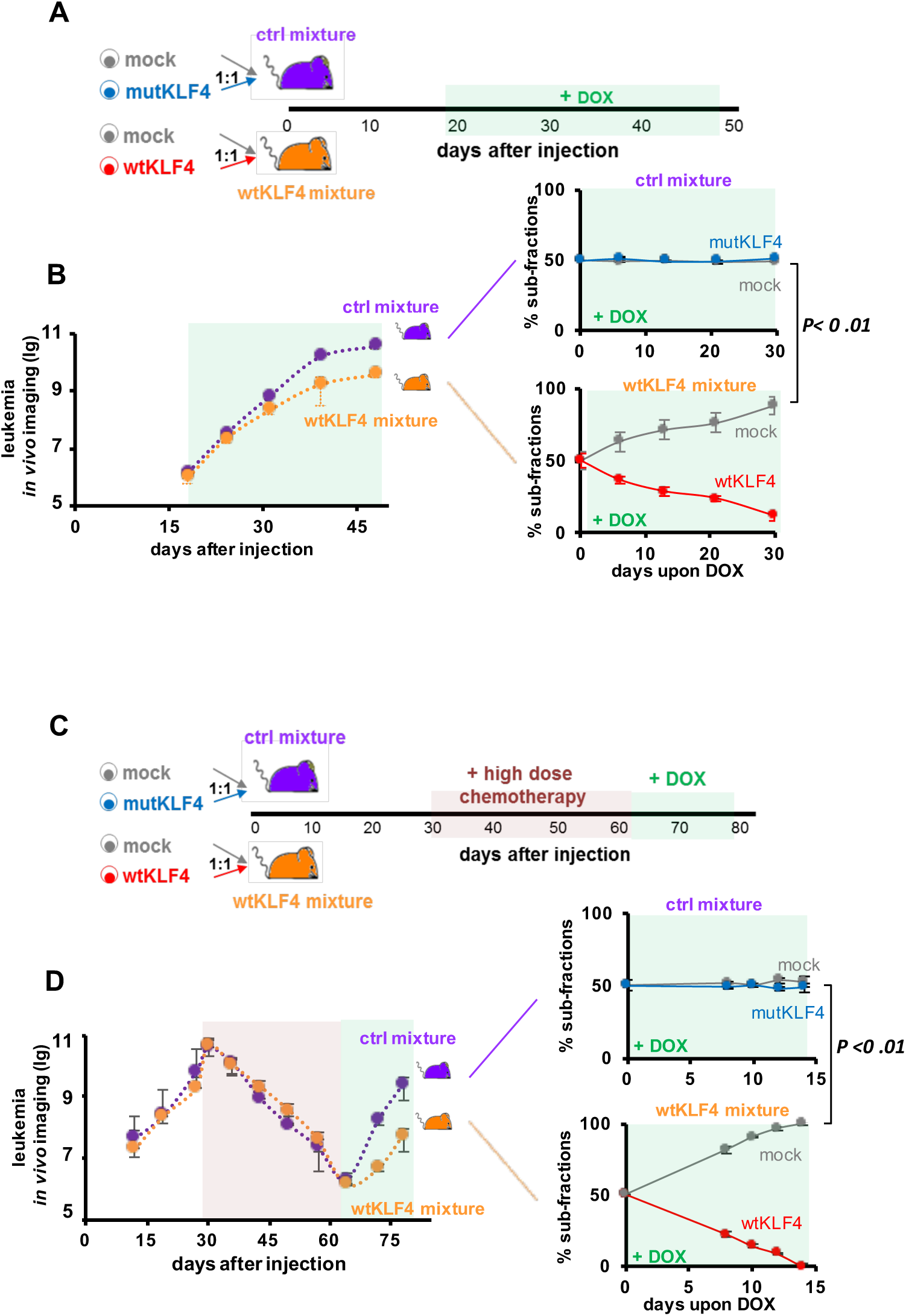
KLF4 re-expressing cells are outcompeted in competitive *in vivo* assays, especially after treatment. **A** A Experimental procedure: NSG mice were injected with 30,000 of the control mixture (purple; n=10) or the wtKLF4 mixture (orange, n=10). The control mixture consisted of a 1:1 ratio of mock and mutKLF4 expressing cells; the wtKLF4 mixture of a 1:1 mixture of mock and wtKLF4 expressing cells. After homing was completed and tumors were established, DOX was added to the drinking water on day 18 and leukemia growth monitored by bioluminescence *in vivo* imaging. 2 mice of each group were sacrificed every week and the proportions of the two sub-fractions in bone marrow analyzed by flow cytometry. **B** Left panel: Leukemia growth as determined by *in vivo* imaging in mice bearing the wtKLF4 or the control mixture. Right panel: sub-fraction analysis by flow cytometry gating on the subpopulation-specific fluorochrome markers T-Sapphire for mock subpopulation and iRFP720 for either mutKLF4 and wtKLF4 subpopulation (see *Supplementary Figure 3B* for constructs). Quantification depicted as mean ± SEM. **p< 0.01 two-tailed Student’s t-test. **C** Experiments were set up as in **A**, C except that DOX was administered after treatment, during tumor re-growth. At high tumor burden (day 31), mice were treated intravenously with high-dose combination chemotherapy (0.25 mg/kg vincristine + 100 mg/kg cyclophosphamide once per week, given on Mondays and Thursdays, respectively) to reduce tumor burden to MRD; at MRD (day 64), chemotherapy was stopped and DOX was added to the drinking water to induce transgene expression. **D** Tumor re-growth after treatment was monitored and analyzed as in **B**.

**Fig. 5.**
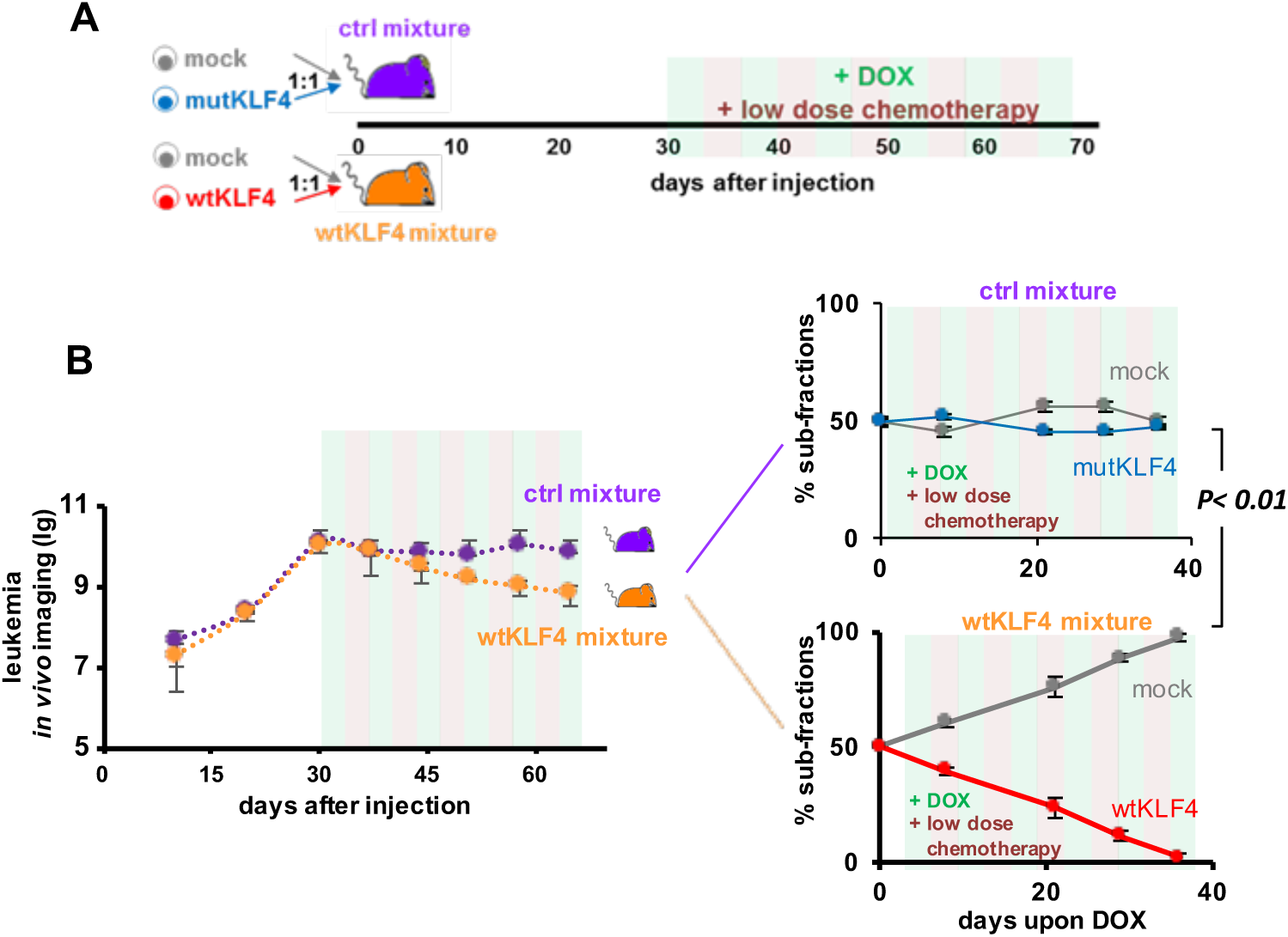
KLF4 expression sensitizes PDX ALL cells to chemotherapy. **A** Experiments were set up as in Figure 4, except that transgene expression was initiated at high tumor burden (day 30) and maintained during low dose combination chemotherapy (0.2 mg/kg Vincristine + 35 mg/kg Cyclophosphamide; days 31-65).**B** Tumor growth was monitored and analyzed as in Figure 4

The effect of KLF4 on tumor growth was further investigated in competition assays *in vivo*, when identical numbers of mock transduced control cells and wtKLF4 transduced cells of the same PDX model were mixed and injected into the same animal (Figure 4A, indicated in orange). As an additional control, a second group of mice was injected with mock and mutKLF4 transduced cells (indicated in purple in Figure 4A). Leukemic load was monitored by *in vivo* imaging and cell ratios of the subpopulations were quantified using the respective fluorochromes.

In these competition experiments, mice injected with the wtKLF4 mixture displayed reduced growth *in vivo*, despite that 50% of cells in the wtKLF4 mixture were control cells (Figure 4B, left panels). Cells transduced with the wtKLF4 allele had a severe disadvantage and nearly disappeared within 30 days (Figure 4B, lower right panel) whereas mock transduced cells expanded readily. In control mice, the ratio of the two subpopulations remained constant over time (Figure 4B, upper right panel) suggesting that genetic manipulation per se did not affect the oncogenic properties of the leukemic PDX models. The competitive *in vivo* assay (Figure 4A and B) confirmed the growth-inhibitory effect of wtKLF4 that we observed in previous experiments (Figure 3). As major advantage, competitive *in vivo* assays can correct for frequent inter-mouse variations, have a high sensitivity and reliability and concomitantly decrease the number of animals in *in vivo* experiments.

Ectopic expression of KLF4 impaired B-ALL growth and survival *in vivo*, but KLF4 might also affect engraftment into the murine bone marrow. To investigate this possibility, we analyzed the homing capacity of KLF4-expressing PDX cells to the murine bone marrow in re-transplantation experiments. mock transduced, wtKLF4 or mutKLF4 expressing PDX cells were isolated from first recipient mice that received DOX at advanced disease stage (*Supplementary Figure 6A*). Equal numbers of cells were combined and re-transplanted into second-generation mice and homing assessed 3 days later (*Supplementary Figure 6A*). The ratio of transplanted mock versus mutKLF4-expressing cells was not altered in the bone marrow after three days, but in reisolated cell populations from mice that had received mock and wtKLF4 transduced cells, the latter were strongly reduced (*Supplementary Figure 6B*), indicating that wtKLF4 expression impairs homing of PDX B-ALL cells *in vivo*. Taken together, re-expression of KLF4 reduced the overall fitness of PDX B-ALL cells *in vivo*. KLF4 functions as a cell cycle inhibitor and pro-apoptotic factor in PDX B-ALL resulting in decreased tumor growth of established tumors and impaired ability to infiltrate the murine bone marrow.

### Treatment-surviving cells are especially sensitive towards re-expressing KLF4

In B-ALL, persistent minimal residual disease (MRD) is the most predictive factors for disease-free survival(11), emphasizing the need to target and eradicate MRD. The number of primary MRD cells isolated from each patient is small by definition, requiring elaborate *in vivo* models to functionally characterize the MRD cell sub-population, a serious technical challenge (39). The inducible expression system provides the means to characterize the role of individual genes at MRD. To mimic the MRD situation in mice, we titrated a combination of the routine drugs Vincristine and Cyclophosphamide at clinically relevant doses to effectively reduce tumor burden by several orders of magnitude over 5 weeks. When MRD was reached (defined as less than 1% leukemia cells in bone marrow) chemotherapy was discontinued and DOX was administered to induce KLF4 expression during the phase of tumor re-growth (Figure 4C).

Treatment response of the control and wtKLF4 mixture was comparable as indicated by *in vivo* imaging (Figure 4D, left panel). In contrast, upon DOX administration at MRD, tumor re-growth was clearly diminished in mice carrying PDX cells transduced with the wtKLF4 mixture (Figure 4D, left panel). Sub-fraction analysis revealed that expression of wtKLF4 (but not mutKLF4) reduced and finally even prevented cellular regrowth *in vivo*, depleting wtKLF4 cells to undetectable levels within two weeks (Figure 4D, right panel). When comparing therapy-naïve, wtKLF4 expressing cells (Figure 4B) with the same cells after chemotherapy at MRD (Figure 4D, lower right panel), KLF4 inhibited growth of MRD cells more drastically than growth of therapy-naïve PDX B-ALL cells (*Supplementary Figure 7*). It thus appears as if MRD cells are especially sensitive towards KLF4 expression suggesting that any regimen that induces KLF4 expression at MRD might be especially effective in tumor consolidation therapy.

### KLF4 sensitizes B-ALL PDX cells towards chemotherapy *in vivo*

Low KLF4 expression levels were a prerequisite for *in vivo* growth of PDX B-ALL samples especially after mice underwent experimental chemotherapy. Our next experiments addressed the question whether re-expression of KLF4 sensitizes patients’ B-ALL cells towards conventional chemotherapy during routine ALL treatment *in vivo*, i.e. in the setting of induction therapy. PDX B-ALL control and wtKLF4 mixtures were injected into mice and DOX was administered together with chemotherapy at high tumor burden (Figure 5A). In this experiment, DOX was administered simultaneously with chemotherapy, which was dosed to stop tumor progression, only, but not reduce tumor size, in order to facilitate identification of KLF4-induced phenotypes.

Accordingly, chemotherapy prevented further tumor progression in mice injected with control cells as intended, but reduced tumor load in mice injected with PDX cells transduced with control and wtKLF4 expressing vectors Figure 5B, left panel). When subcellular fractions of these animals were analyzed, the number of wtKLF4 but not mutKLF4 expressing cells was significantly decreased by chemotherapy such that wtKLF4 expressing cells were outcompeted and were lost within less than 40 days (Figure 5B, right panel). KLF4 re-expression also sensitized NALM-6 cells towards chemotherapy *in vitro* (*Supplementary Figure 8*) confirming our data with PDX B-ALL cells *in vivo*. The data suggest that upregulation of KLF4 may synergize with standard therapeutic regimens, i.e. conventional chemotherapy to eliminate B-ALL cells in patients.

### Azacitidine-induced cell death partially depends on KLF4

Next, we searched for drugs capable of upregulating KLF4. We first tested the small molecule APTO-253, which was introduced as a KLF4 inducing drug (45), but was recently shown to also regulate MYC (46). APTO-253 indeed upregulated KLF4 in our hands and sensitized cell lines towards Vincristine treatment *in vitro* (*Supplementary Figure 9*), indicating that KLF4 can be re-upregulated by drugs in B-ALL, in principle. We speculated that existing approved drugs might be able to re-upregulate KLF4 in B-ALL to facilitate clinical translation. The demethylating agent 5-Azacitidine (Aza) was shown to upregulate KLF4 expression in other tumor entities (23, 27, 30, 47). Although Aza can reverse DNA hypermethylation, it regulates protein expression on multiple levels, and its decisive mode of action as an anti-cancer agent remains, at least in part, elusive (48). Aza is increasingly used in clinical trials on patients with hematopoietic malignancies and during MRD, e.g., in acute myeloid leukemia, to prevent or retard relapse (49). Here we found that Aza upregulated KLF4 levels in B-ALL PDX cells as well as in B-ALL cell lines (Figure 6A and *Supplementary Figure 10A*), and decreased cell viability *in vitro* at clinically relevant doses (*Supplementary Figure 10A*). To assess whether Aza-induced cell death in B-cell tumor cells is mediated by increased levels of endogenous KLF4 protein, we generated KLF4 knockout (KO) cells, which is a challenging step in PDX. For B-ALL PDX, cells were transduced with a lentiviral vector to introduce Cas9 (Figure 6B and *Supplementary Figure 10B*). In subsequent steps, a second vector encoding a single KLF4 targeting guide RNA and a reporter construct to enrich successfully gene-edited cells were introduced. PDX B-ALL cells with stable KO of KLF4 could readily be established (Figure 6C). Stimulation of PDX B-ALL cells with Aza at clinically relevant concentrations reduced cell viability in control cells but not in PDX B-ALL or NALM-6 cells with KLF4 KO (Figure 6D and *Supplementary Figure 10C*). These data suggest that KLF4 at least partially mediates the effect of Aza in B-ALL cells.

**Fig. 6.**
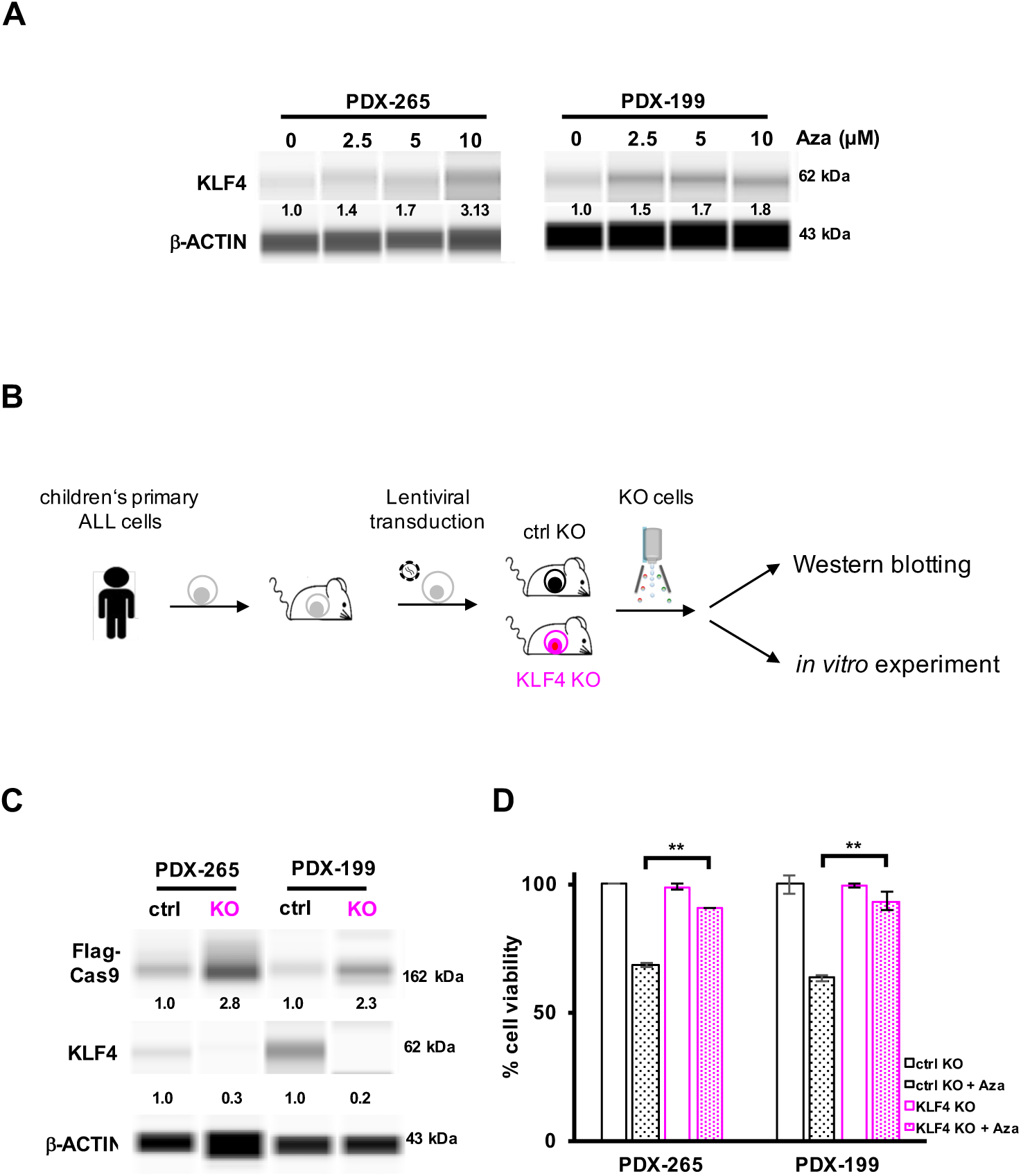
Azacitidine-induced cell death depends on KLF4. **A** PDX-265 and PDX-199 cells were treated with different concentrations of Azacitidine (Aza) for 48h. KLF4 protein expression was analyzed by capillary immunoassay, *β*-actin served as loading control. One representative analysis of 2 independent experiments is shown. **B** Experimental procedure: B-ALL PDX cells were lentivirally transduced with Cas9, sgRNA and reporter expression vectors (*Supplementary Figure 10B*) and injected into NSG mice to generate ctrl KO (black) and KLF4 KO (pink) PDX cells. Mice were sacrificed at full blown leukemia and marker-positive populations enriched by flow cytometry, gating on the recombinant markers (mTaqBFP for Cas9, mCherry for the sgRNAs and GFP for the reporter construct) and subjected to capillary immunoassay and *in vitro* culture. **C** KLF4 protein level of ctrl KO or KLF4 KO PDX-265 and PDX-199 cells were analyzed by capillary immunoassay. Due to low expression of KLF4 in the ctrl cells, images are displayed with strongly increased contrast; *β*-actin served as loading control. One representative analysis out of 2 experiments is shown. **D** PDX B-ALL ctrl KO cells and KLF4 KO cells were treated with 2.5 µM Aza *in vitro* for 48h and cell viability was measured by flow cytometry. Viability was normalized to non-treated cells. Mean ± SEM of duplicates is shown. ** p<0.01 by two-tailed unpaired t test.

Taken together, our data show that KLF4 might represent a promising novel therapeutic target for B-ALL. Our data also document that Aza, an established drug, can upregulate KLF4. Introducing Aza into standard poly-chemotherapy protocols of B-ALL patients with the intention to raise KLF4 levels might reduce tumor burden and increase sensitivity towards conventional chemotherapy. Patients with B-ALL might benefit from this option that needs to be tested for proof of concept in clinical trials.

## Discussion

We describe a novel method that allows characterizing which alterations in tumor cells harbor functional relevance, in preclinical PDX models *in vivo*. Using the technique, we demonstrate that PDX B-ALL models depend on downregulation of KLF4; re-upregulating KLF4 using the inducible expression system or by treatment with Aza impairs tumor maintenance *in vivo*. We work with genetically engineered PDX models that we propose calling GEPDX models, in accordance to genetically engineered mouse models (GEMM). Inducible GEMM were designed to allow determining the role of single molecules independently from gestation, for example. In analogy, our novel inducible GEPDX models allow determining the role of single molecules in pre-established tumors, independently from, e.g., tumor transplantation into mice. Inducible GEPDX models might be established in a broad range of different tumor types beyond B-ALL and hold the potential to interrogate gene function in preclinical cancer models at clinically relevant time points, such as at MRD. Using these approaches, we demonstrate that KLF4 i) limits B-cell transformation by EBV ii) impairs B-ALL maintenance *in vivo* iii) strongly reduces regrowth of B-ALL PDX cells in the situation of MRD and; iv) increases response of PDX B-ALL cells to therapy. We show that the well-described roles of KLF4 as a quiescence factor and apoptosis-inducer are maintained in the malignant B-cells studied (13). Of major translational importance, we show that Aza upregulates KLF4 in B-ALL and upregulated KLF4 mediates, at least in part, the anti-leukemia effect of Aza.

Aza has previously been shown to upregulate KLF4 in other tumor entities (23, 30, 50–52) and we demonstrate that this ability is maintained in B-ALL. A proven clinical benefit of Aza treatment in hematological malignancies was first obtained in myelodysplastic syndrome and AML (48), without increasing rates in secondary malignancies. In B-ALL, case reports exist on the clinical activity of Aza (53–56) and an ongoing clinical study applies Aza in patients with KMT2A re-arranged B-ALL (NCT02828358). As molecular mechanism of Aza-mediated antitumor activity remain incompletely defined (48), our data suggests that KLF4 upregulation represents an important mechanism of Aza to eliminate B-ALL cells.

The data presented here highlight the power of molecular / functional testing to improve understanding of alterations in individual tumors and link gene function to available drugs. As we discovered that downregulating KLF4 plays an essential role for established PDX models of B-ALL to grow *in vivo*, and the widely used drug Aza upregulates KLF4, our data strengthen the idea of applying Aza in patients with B-ALL.

## Materials and Methods

### Ethical Statements

For the two primary B-ALL patient samples, written informed consent was obtained from all patients or from parents/caregivers in cases when patients were minors. The study was performed in accordance with the ethical standards of the responsible committee on human experimentation (written approval by Ethikkommission des Klinikums der Ludwig-Maximilians-Universität Munich, number 068-08 and 222-10) and with the Helsinki Declaration of 1975, as revised in 2000. Animal trials were performed in accordance with the current ethical standards of the official committee on animal experimentation (written approval by Regierung von Oberbayern; July 2010, number 55.2-1-54-2531-95-10; July 2010, 55.2-1-54-2531.6-10-10; January 2016, ROB-55.2Vet-2532.Vet02-15-193; May 2016, ROB-55.2Vet-2532.Vet02-16-7 and August 2016, ROB-55.2Vet-2532.Vet03-16-56).

### Genetic engineering in EBV

In the maxi-Epstein Barr virus (EBV) plasmid, wtKLF4 and mutKLF4 cDNAs were fused to the 3’ open reading frame of the viral EBNA2 by a T2A element, mediating co-expression of both genes from the same promoter. While the wtKLF4 construct contained the entire open reading frame, the mutKLF4 construct lacked the two N-terminal zinc finger domains (57).

### Genetic engineering of PDX B-ALL cells for inducible transgene expression

Primary patients’ B-ALL cells were transplanted into immunocompromised mice to generate PDX models. PDX B-ALL were lentivirally transduced and transgenic cells enriched using flow cytometry gating on recombinant fluorochromes as described (58). For inducible transgene expression, PDX B-ALL cells were lentivirally transduced with three consecutive constructs containing the tet activator, the tet repressor and KLF4 expression cassettes under control of the TRE promoter (44).

### *In vivo* experiments

Leukemia growth and treatment effects were monitored using bioluminescence *in vivo* imaging as described (58). Competitive experiments were performed by mixing two derivate cell populations, each expressing a different transgene and distinct fluorochrome marker, and injecting both into the same animal. Human PDX cells were isolated and enriched from murine bone marrow or spleen as described (39) and distribution of each subpopulation measured by flow cytometry using the different recombinant fluorochrome markers.

### Methods detailed in the supplement

Details are provided for constructs for inducible expression of KLF4 and CRISPR/Cas9-mediated knockout, cell isolation and culture conditions, EBV infection, gene expression analysis, *in vitro* culture, *in vivo* growth of PDX-ALL, monitoring of tumor burden and drug treatments, homing assay, *in vitro* culture of PDX cells and drug treatment, cell cycle analysis, gene expression analysis, capillary immunoassay and statistical analysis.

## Supporting information

Liu et al_Supplemental Information

## ACKNOWLEDGEMENTS

We thank Daniela Senft for drafting the manuscript and valuable discussions; Liliana Mura, Fabian Klein, Maike Fritschle, Annette Frank and Miriam Krekel for excellent technical assistance; Markus Brielmeier and team (Research Unit Comparative Medicine) for animal care services; We thank Jean Pierre Bourquin and Beat Bornhäuser for providing engrafted sample ALL-265.

