## Supplementary material for "Inducible transgene expression in PDX models *in vivo* identifies KLF4 as therapeutic target for B-ALL": Liu et al_Supplemental Information

<u>This file contains:</u>

Supplementary Methods

Supplementary References

Supplementary Tables

Supplementary Figures and Legends

### Supplementary Methods

#### Constructs for inducible KLF4 expression and KLF4 knockout

Tet-on inducible expression system (1): The pRRL-SFFV-rtTA3-IRES-Luc2-P2A-mtagBFP was a gift from Johannes Zuber (Research Institute of Molecular Pathology, Vienna, Austria). The two DNA fragments that contained the coding sequences of tetracycline repressor protein (tetR), coupled via T2A either to iRPF720 or T-Sapphire were synthesized (IDT DNA technologies, Coralville, IA), and cloned into the pCDH vector (Addgene plasmid #104834) using BamHI and SalI. For KLF4 transgene expression, The wtKLF4-T2A-mCherry and mutKLF4-T2A-mCherry DNA fragments were synthetized (IDT DNA technologies), amplified, and cloned into the pRRL-TRE-mCherry-PGK-Venus vector (based on TRMPV, Addgene plasmid # 27991, but dsRED was replaced by mCherry) using BamHI and XhoI. mutKLF4 lacks the amino acids 462-513 (NP_001300981.1), encoding zinc finger domain 2 and 3 (2).

CRISPR/Cas9 expression vectors: A DNA fragment containing the cDNA of spleen focus forming virus (SFFV) promoter (Addgene plasmid #27348), humanized Staphyococcus pyogenes Cas9 (hSpCas9 PX330, Addgene plasmid #42230) and a T2A-linked mTaqBFP fluorochrome (pTagBFP-C vector, PF171, Envrogen, Lawrenceville, NJ) was amplified by PCR and cloned into the pCDH vector (Addgene Plasmid #72263) using the SpeI/EcoRI, EcoRI/BamHI and NsiI/SalI restriction sites, respectively. For sgRNA expression, the pCDH vector (Addgene Plasmid #104833) expressing mCherry under control of the Elongation factor 1-alpha (EF1$\alpha$) promoter was complemented by an additional H1 promoter expressing the sgRNA; towards this aim, an oligonucleotide containing the H1 promoter, a BbsI cutting site (Addgene plasmid #42230), a gRNA scaffold sequence (Addgene plasmid #42230) and a polymerase III terminator sequence (gift from Macro J. Herold, Walter and Eliza Hall Institute, Australia) were ordered from IDT DNA technologies and ligated into the pCDH backbone between the cPPT and EF1$\alpha$ promoter using ClaI and SpeI. The sgRNA sequences (Table S2) were synthesized (Eurofins MWG-Operon, Ebersberg, Germany), annealed and ligated into the vector. A detailed description of the CRISPR/Cas9 reporter system used will be provided upon request.

**EBV-induced transformation of normal naïve human B-cells**

EBV constructs: The wild-type EBV strain, termed r_wt/B95.8 (6008) (3) is based on the maxi-EBV plasmid p2089, which comprises the entire B95.8 EBV genome cloned onto a mini-F-factor plasmid in *E. coli* (4). The EBV construct carries the right handed oriLyt and expresses all 25 EBV-encoded pri-miRNAs from their endogenous viral promoters as well as the LF1, LF2 and LF3 genes that are absent in B95.8 DNA reconstructing EBV field strains.

Two mutant EBV strains that express the *KLF4* cDNA (6948_EBNA2-T2A-KLF4; wtKLF4) and a DNA-binding deficient mutant KLF4 cDNA (6949_EBNA2-T2A-mutKLF4; mutKLF4 (2)) were constructed (Supplementary Figure 2C). wtKLF4 and mutKLF4 cDNAs were fused to the 3' open reading frame of the viral EBNA2 by a T2A element, mediating co-expression of both genes from the same promoter (Supplementary Figure 2C). Details of the genetic engineering of the two mutant EBVs and their DNA sequences are available upon request.

Isolation of naïve B-lymphocytes: Adenoid biopsies from children were mechanically disintegrated to obtain single cell suspensions. Human B lymphocytes were separated from T cells by rosetting with defibrinated sheep blood and purified by Ficoll–Hypaque (Biochrom AG, Berlin, Germany) density gradients centrifugation. The interphase with the B lymphocytes was removed and washed three times with PBS. The cells were counted and directly used for staining with antibodies against CD38 (#25-0389-42, eBioscience, San Diego, USA) and IgD (#555778, BD Pharmingen, San Diego, USA). Naïve B-lymphocytes ($IgD^+/IgH^+$ and $CD38^-$) were physically sorted with a BD FACSAria™ III instrument (Becton-Dickinson, San Jose, USA) and used for further experiments.

EBV infection and *in vitro* culture: Primary B cells were infected with mock-EBV (r_wt/B95.8 (6008)), wtKLF4-EBV (6948), and mutKLF4-EBV (6949) with a multiplicity of infection (MOI) of 0.1 overnight. The next day, the medium was replaced with fresh medium and the infected cells were seeded in 96-well cluster plates at an initial concentration of $10^6$ cells/ml.

#### Cell isolation and *in vitro* culture conditions

Cell culture conditions: B-cells were cultivated in RPMI1640 medium supplemented with 8% FCS, 100 U/ml streptomycin/penicillin, 1 mM sodium pyruvate, 100 mM sodium selenite, and 0.43% α-thioglycerols. NALM-6 (human pre B-ALL) and RAJI (Burkitt lymphoma cell line, EBV-positive) were purchased from DSMZ (German collection of microorganisms and cells, Braunschweig, Germany) and were maintained in RPMI-1640 supplemented with 2 mM L-glutamine (Life Technologies GmbH, Darmstadt, Germany) and 10% FCS (Biochrom AG). HEK293T cells were maintained in DMEM (Life Technologies) supplemented with 2 mM L-glutamine and 10% FCS. PDX cells were cultured in RPMI medium supplemented with 20% FSC, 1% pen/strep, 1% gentamycin, 6 mg/l insulin, 3 mg/l transferrin, 4 µg/l selenium (Gibco, San Diego, USA), 2 mM glutamine, 1 mM sodium pyruvate, 50 µM α-thioglycerol (Sigma-Aldrich, St. Louis, USA).

Peripheral blood mononuclear cells (PBMCs) were isolated on Ficoll–Hypaque density gradients (Biochrom AG) from 5ml of heparinized venous blood drawn from healthy volunteer donors.

#### Lentiviral transduction and enrichment of PDX ALL cells

PDX ALL-265 and ALL-199 were established as described (5); Lentiviruses were produced and PDX cells genetically engineered as described (6,7) and were titrated such that transduction efficiencies were below 1%, to achieve a single genomic integration per cell. As unique variance and to save resources, lentivirally transduced PDX cells were kept in culture to await fluorochrome marker expression; on day 4-6, successfully transduced cells were enriched by flow cytometry using a BD FACSAria™ III cell sorter (Becton-Dickinson) and injected into mice for amplification, as highly transgenic population.

Isolation of transgene-expressing cell populations from murine bone marrow or spleen was performed as described (5). Based on fluorochrome markers, viable cells were either enriched for further experiments or analyzed using the LSRII flow cytometer (Becton-Dickinson).

**PDX *in vivo* experiments**

PDX *in vivo* experiments were performed and monitored by bioluminescence *in vivo* imaging as previously described (5). To induce KLF4 transgene expression, Doxycyline (DOX) was solved at 1 mg/ml in drinking water containing 5% Glucose (Sigma-Aldrich) and served to mice at free disposition (wtKLF4 or mutKLF4); DOX-containing drinking water was changed weekly.

For pairwise competitive *in vivo* experiments, PDX-ALL cells expressing mock and wtKLF4 (wtKLF4 mixture) or mock and mutKLF4 (control mixture) were mixed at a ratio of 1:1 and injected into the tail vein of NSG mice.

*In vivo* chemotherapy treatment: PDX ALL bearing mice were treated with PBS or a combination of vincristine (VCR; Merck, Darmstadt, Germany; given once per week on Fridays) and cyclophosphamide (Cyclo; Baxter, USA; given once per week on Mondays). In low dose regimens, mice received 0.2 mg/kg VCR plus 35 mg/kg Cyclo until the experimental endpoint; in high dose regimens, mice received 0.25 mg/kg VCR + 80 mg/kg Cyclo.

*In vivo* homing assay**:** Mice were injected with PDX-265 cells transduced with mock, mutKLF4 or wtKLF4 constructs; at day 30, when tumor burden was high, DOX was administered to induce transgene expression. 2 days later, PDX cells were harvested from the spleen and mCherry-positive cells were sorted using BD FACSAria$^{TM}$ III. $5x10^5$ PDX-265 cells each were mixed at 1:1 ratio and reinjected into NSG recipients. 3 days after injection, mice were sacrificed and bone marrow cells were isolated. PDX-265 cells were enriched by Mouse cell depletion kit (Miltenyi Biotec, Bergisch Gladbach, Germany) as previously described (7) and the percentage of mock and wtKLF4/mutKLF4 analyzed by flow cytometry.

***In vitro* drug treatment**

For *in vitro* drug treatment, PDX cells or NALM-6 cells were cultured and dilutions in PBS from either Vincristine (1mg/ml stock), APTO-253 (10 mM stock in DMSO, HY-16291, MedChem Express, New Jersey, USA) or Azacitidine (10 mM stock in DMSO, Biozol Diagnostica, Eching, Germany) added to the medium and cells were analyzed 2 days after treatment.

#### Analysis of cell viability and apoptosis

Cell viability was determined by flow cytometry based on FSC/SSC gating and/or staining with Annexin V. Viability of EBV-infected B cells was measured as previously described (8). In brief, EBV-infected cells were washed twice with FACS staining buffer (PBS, 0.5 % BSA, 2 mM EDTA), resuspended in 200 µl Annexin V staining buffer together with 2 µl Cy5-coupled Annexin V (BioVision, Milpitas, USA) and incubated in the dark for 10 min. Percentage of Annexin V-positive cells was determined using a BD FACS Fortessa instrument (Becton-Dickinson).

#### Cell cycle analysis

BrdU staining: Per time point, $5x10^5$ EBV-infected cells were seeded initially and cultivated further. Prior to harvest, the cells were incubated with BrdU (10 µM final concentration) and incubated in the dark at 37°C for 1 hour. After BrdU incorporation, the cells were washed with FACS staining buffer and resuspended in Cytofix/Cytoperm (Becton-Dickinson) buffer followed by a 15-20 min incubation on ice. The cells were washed with FACS staining buffer and frozen in freezing medium (90 % FCS, 10 % DMSO). The samples were stored at -80°C for further analysis. The BrdU incorporation was analyzed using the APC BrdU Flow Kit (#552598, BD Pharmingen) according to the manufacturer's recommendations. The cells were thawed, re-fixated with BD Cytofix/Cytoperm buffer for 5 min on ice, washed with BD Perm/Wash buffer, and resuspended in PBS and DNase. The mix was incubated at 37°C for 1 h, then washed with BD Perm/Wash buffer, resuspended in BD Perm/Wash buffer and stained with 1 µl anti-BrdU-APC coupled antibody for 20 min at RT. Afterwards, the cells were washed with BD Perm/Wash buffer, resuspended in 7-AAD solution (20 µl per sample of a 50 µg/ml stock solution, (#420404 Biolegend, San Diego, USA) and FACS staining buffer. After 10 min incubation, the samples were measured with a BD FACS Fortessa instrument and analyzed with the FlowJo software (Tree Star Software, Ashland, USA) to determine the cell cycle distribution.

Propidium Iodide (PI) staining

$10^6$ viable, mCherry positive sorted cells were fixed in 70% ethanol and stored at -20 $^o$C overnight. The next day, cells were washed in cold PBS, stained with 50 µg/ml PI

(Sigma-Aldrich) and incubated for 30 min at 37 °C. Afterwards, cells were measured by flow cytometry and analysis was performed using the FlowJo software (Tree Star Software).

#### Gene expression analysis

Array based expression analysis: The data set of B-ALL (n=306) was used for gene expression analysis (GSE66006, GSE78132, GSE37642) (9-11). Details of the diagnostic work-up, sample preparation, hybridization, image acquisition and analysis have been described previously (9). In brief, for probes to probe set annotation, we used custom chip definition files (12). The HG-U133 A, B chips and HG-U133 Plus 2.0 chips were normalized separately by the robust multichip average method as described (13) and only probes present on all chips were included in the analysis (n=17,389). The batch effect resulting from the use of different chip designs was corrected by applying an empirical Bayesian method as described elsewhere (14).

RNA sequencing and data analysis of primary ALL patient samples at diagnosis and 33 days following therapy was described previously; at MRD, the analyzed cells consisted of highly enriched ALL cells, distinguished from normal bone marrow cells by individualized immune surface markers (5).

RNA-seq based gene expression analysis at early time points following EBV infection: A detailed description of the time course experiment has recently been published by Mrozek-Gorska et al. (15). Raw data are available at the following link (https://www.ebi.ac.uk/arrayexpress/experiments/E-MTAB-7805/) and processed data can be freely accessed online (http://ebv-b.helmholtz-muenchen.de/).

RT-PCR: Total RNA was extracted from $10^6$ cells using the Qiagen RNeasy Kit (Hilden, Germany) according to manufacturer's instructions. 100 ng RNA was reverse transcribed using iScript cDNA (Bio-Rad). cDNA and primers (Table S2, Eurofins MWG-Operon) were mixed with PCR Master Mix (Thermo Scientific , Waltham, USA) and amplified in 25 cycles (30 sec at 94°C, 30 sec at 60°C, 30 sec at 72°C). Amplified cDNA was loaded on a 2% agarose gel (Invitrogen, Carlsbad, USA) and bands were visualized following staining with Midori$^{Green}$ Advance (Nippon Genetics, Düren,

Germany) using the gel imaging system E Box CX5 (Vilber Lourmat, Marne La Vallée, France).

##### Protein expression analysis

Flow cytometry-enriched cell populations were incubated in lysis buffer (#9803, Cell Signaling Technology, Boston, USA) on ice for 30 min. Protein concentration was measured by BCA assay (#7780, New England Biolabs, Beverly, USA) and abundance of specific proteins determined by two different methods.

1. conventional Western blotting

Equal amounts of protein were separated using 12% gels and transferred to a nitrocellulose membrane (TransBlott, Bio-Rad, Munich, Germany), incubated with antibodies against Caspase3 or PARP, washed and incubated with a secondary antibody coupled to an enzyme (GE Healthcare, Menlo Park, USA). Enyzme activity was visualized using a chemiluminescent substrate (Thermo scientific, USA).

2. capillary protein immunoassay

The technology (WES, Simple Western, ProteinSimple, San Jose, USA) enables a Western-Blot-like analysis with the advantage of requiring only low cell numbers, typically 10.000 cells per lane. It was used as only minor cell numbers can be re-isolated from each mouse.

Procedures were performed following manufacturer's instructions; in brief, each capillary was loaded with protein lysate and electrophoresis was performed; capillaries were processed to attach all proteins to the capillary wall and incubated with a single antibody; results were measured as emission curves from each capillary and "Western-Blot-like presentations" calculated thereof using the Compass software (ProteinSimple), including quantification. Final "Western-Blot-like presentations" appear clearly different from conventional Western Blots; due to very high sensitivity, equal loading is hard to achieve, but also not required, as protein amounts are calculated with high sensitivity and reliability.

Primary antibody against KLF4 was purchased from R&D systems (AF3640, Minneapolis, MN, USA), against PARP from Cell Signaling Technology (9542T, USA), against GAPDH from R&D systems (MAB5718-SP, USA), against Caspase-3 from Millipore (05-654, Billerica, USA) and against β-ACTIN from Novus biologicals (NB600-501SS, Littleton, USA).

**Statistical Analysis**

Statistical significance was determined using the “R” software package. Mean values were compared by Student's t-test or analysis of variance, depending on the number of groups and days to be analyzed.

### Supplemental Tables

**Table S1 sgRNA sequences**

| | sgRNA sequence |
|---|---|
| *Gaussia luciferase (GLuc)* | GGACTCTTTGTCGCCTTCGT |
| KLF4 seq#1 | GTGGTGGCGCCCTACAACGG |
| KLF4 seq#2 | GAGCGATACTCACGTTATTCG |
| KLF4 seq#3 | GTCGGTCATCAGCGTCAGCAA |

**Table S2 RT-PCR primer sequence**

| name | Primer sequences |
|---|---|
| GAPDH | forward: GAAGGTGAAGGTCGGAGTC |
| | reverse: GAAGATGGTGATGGGATTTC |
| KLF4 | forward: CGGGCTGATGGGCAAGTT |
| | reverse: GGGCAGGAAGGATGGGTAA |

### Supplementary Figures

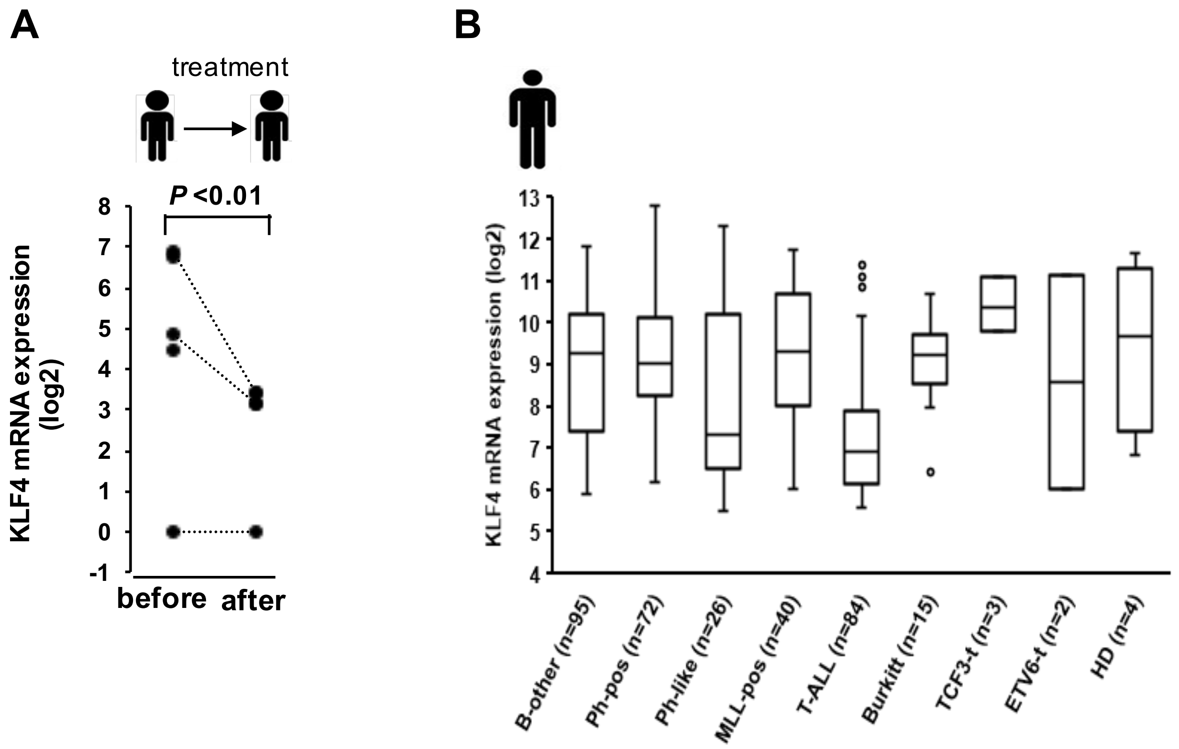


**Figure S1 KLF4 mRNA is downregulated in primary patient leukemia cells**

A. Expression of KLF4 mRNA in primary samples from pediatric ALL patients at diagnosis (n=5, before) and after the first block of treatment according to the BFM-2009 protocol (n=3, after, day 33). For details on cell sorting and RNA sequencing see Ebinger et al. 2016; RNA-seq data are publicly available (GSE83142). Dashed lines indicate matched samples; p<0.01 by two-tailed unpaired t test.

B. Expression of KLF4 mRNA in bone marrow from 306 ALL patients. Datasets are publicly available (GSE66006, GSE78132, GSE37642). Ph-pos: Philadelphia-positive; HD: Hyperdiploidy.

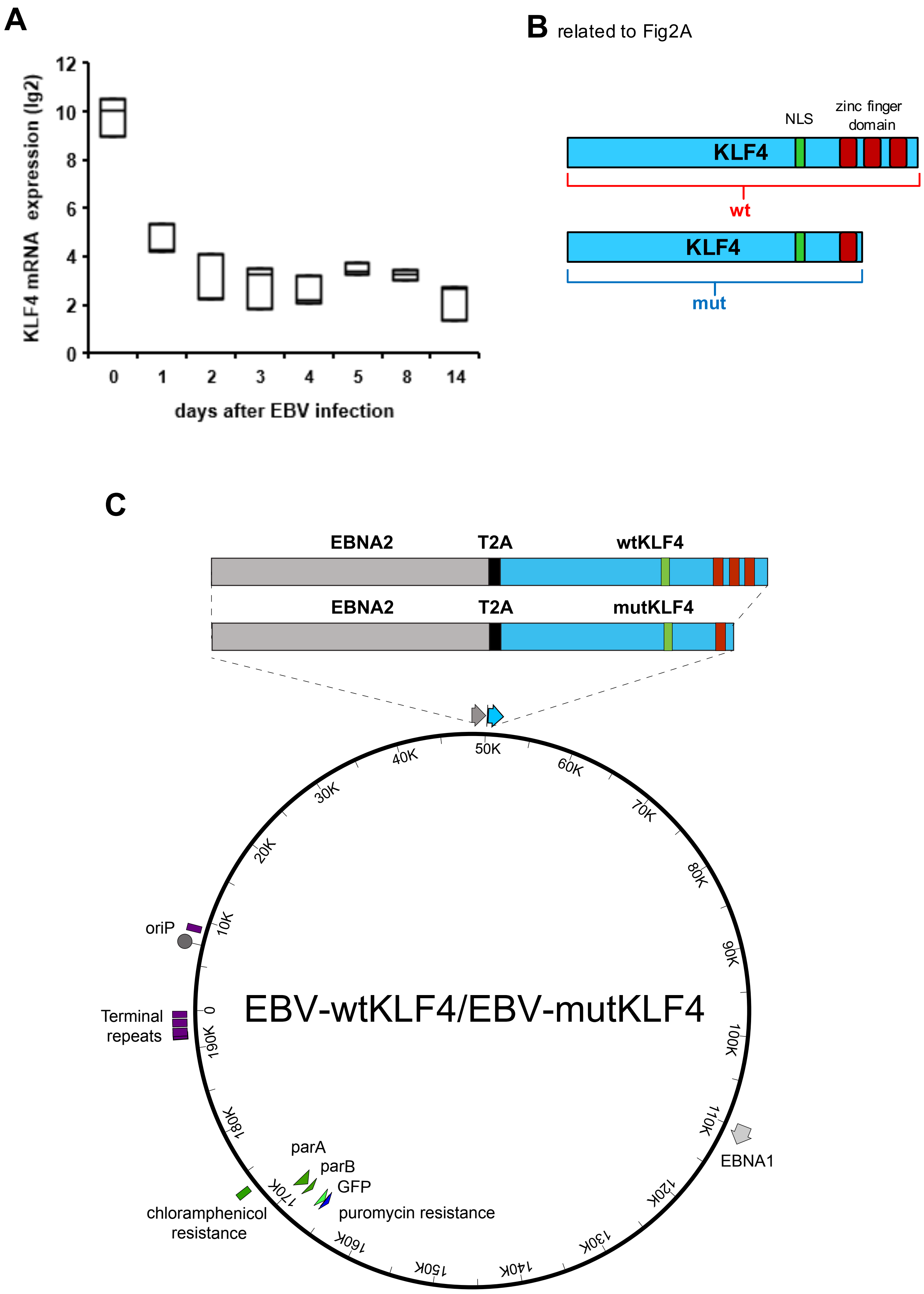
A
KLF4 mRNA expression (lg2)
12
10
8
6
4
2
0
0
1
2
3
4
5
8
14
days after EBV infection
B related to Fig2A
NLS
zinc finger domain
KLF4
wt
KLF4
mut
C
EBNA2
T2A
wtKLF4
EBNA2
T2A
mutKLF4
EBV-wtKLF4/EBV-mutKLF4
oriP
Terminal repeats
EBNA1
parA
parB
GFP
puromycin resistance
chloramphenicol resistance

**Figure S2 KLF4 mRNA expression and KLF4 expression constructs**

A. Primary normal B-cells were infected with EBV as in printed Figure 1 and kinetic extended to 14 days; KLF4 mRNA was measured at the time points indicated as described in Mrozek-Gorska et al. 2019; mean ± SD of 3 independent experiments is shown.

B. Scheme of KLF4: At the carboxy terminus, the KLF4 protein contains 3 zink finger domains (red blocks) which are responsible for DNA binding at GC-rich sequences; the nuclear localization signal (NLS) is indicated by a green box. A truncated mutKLF4 version was generated lacking the two of the three C-terminal zinc finger domains.

C. Scheme of the EBV genome modified to express wtKLF4 or mutKLF4. KLF4 cDNA was linked to the viral EBNA2 sequence via the self-cleaving 2A sequence (T2A); expression of either wtKLF4 or mutKLF4 was driven from the endogenous Wp promoter which also drives EBNA2.

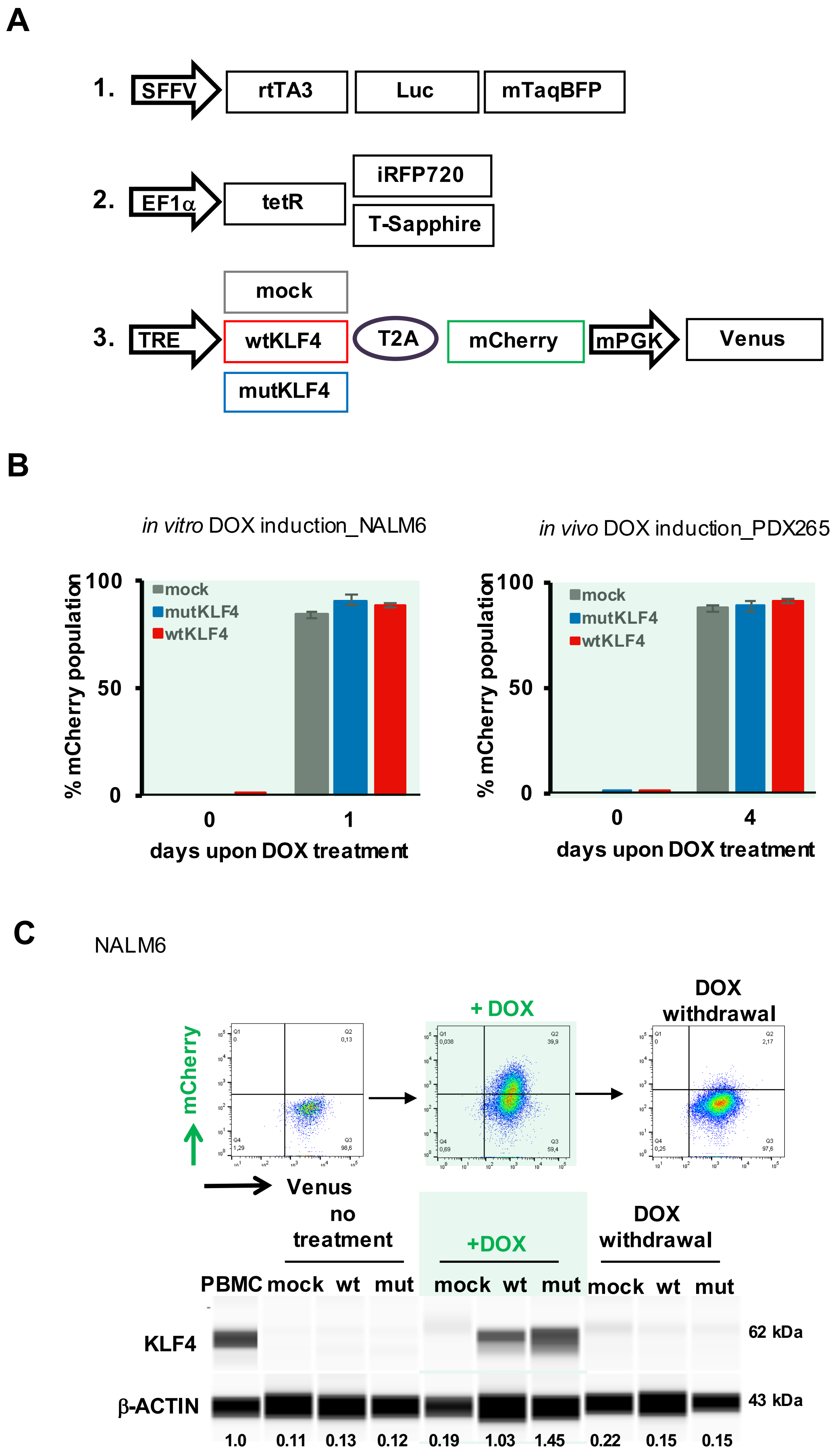

A
1.
SFFV
rtTA3
Luc
mTaqBFP
2.
EF1α
tetR
iRFP720
T-Sapphire
3.
TRE
mock
wtKLF4
mutKLF4
T2A
mCherry
mPGK
Venus
B
in vitro DOX induction_NALM6
in vivo DOX induction_PDX265
% mCherry population
100
50
0
mock
mutKLF4
wtKLF4
0
1
4
days upon DOX treatment
C
NALM6
+ DOX
DOX withdrawal
mCherry
Venus
no treatment
+DOX
DOX withdrawal
PBMC
mock
wt
mut
KLF4
β-ACTIN
62 kDa
43 kDa
1.0
0.11
0.13
0.12
0.19
1.03
1.45
0.22
0.15
0.15

**Figure S3 A tet-on inducible system to re-express KLF4 in PDX ALL cells**

A. Scheme of expression vectors: To enable inducible expression, three constructs were used.

1. The first construct expressed the reverse tet-responsive transactivator 3 (rtTA3) together with Luciferase (Luc) for bioluminescence *in vivo* imaging, and mTaqBFP for enriching transgenic cells by flow cytometry.

2. The second construct expressed the tet repressor (tetR) to reduce leakiness together with either one of two different fluorochromes (iRFP720 or T-Sapphire) to distinguish two different cell populations in a single mouse during competitive *in vivo* assays.

3. The third construct contained either mock, wtKLF4 or mutKLF4, expressed under control of the DOX-inducible tetracycline response element (TRE); a second fluorochrome (mCherry) was directly coupled to KLF4 by a T2A linker sequence, enabling visualization and flow cytometric enrichment of cells successfully expressing the induced transgene upon DOX treatment; in addition, the third expression construct constitutively expressed the fluorochrome Venus for enrichment of transduced cells.

B. High expression levels of mCherry after DOX treatment.
mCherry expression construct is linked to the expression construct of wtKLF4 or mutKLF4 by a T2A site such that mCherry indicates expression of KLF4.

1. Left: $10^6$ NALM-6 cells transduced with the indicated constructs were incubated with 0.5µg/ml DOX for 24h and the percentage of mCherry positive cells was determined by flow cytometry.

2. Right: PDX ALL-265 cells transduced with the indicated constructs, enriched by flow cytometry and 60,000 cells injected into NSG mice. DOX was added to the drinking water of mice upon high tumor burden. After 4 days of DOX treatment, mice were sacrificed and percentage of mCherry positive cells from spleen and bone marrow determined by flow cytometry.

C. Reversibility of the tet-on expression system. NALM-6 cells were transduced with mock, wtKLF4 or mutKLF4 constructs and cultured without or with 0.5µg/ml DOX overnight. Cells were either harvested, or DOX withdrawn and cells cultured for additional 5 days in the absence of DOX. Upper panel: Representative dot plots of NALM-6 cells transduced with the mock construct; expression of mCherry is linked to expression of KLF4 and induced by DOX. Lower panel: KLF4 protein expression was analyzed by capillary immunoassay; beta-actin was used as loading control.

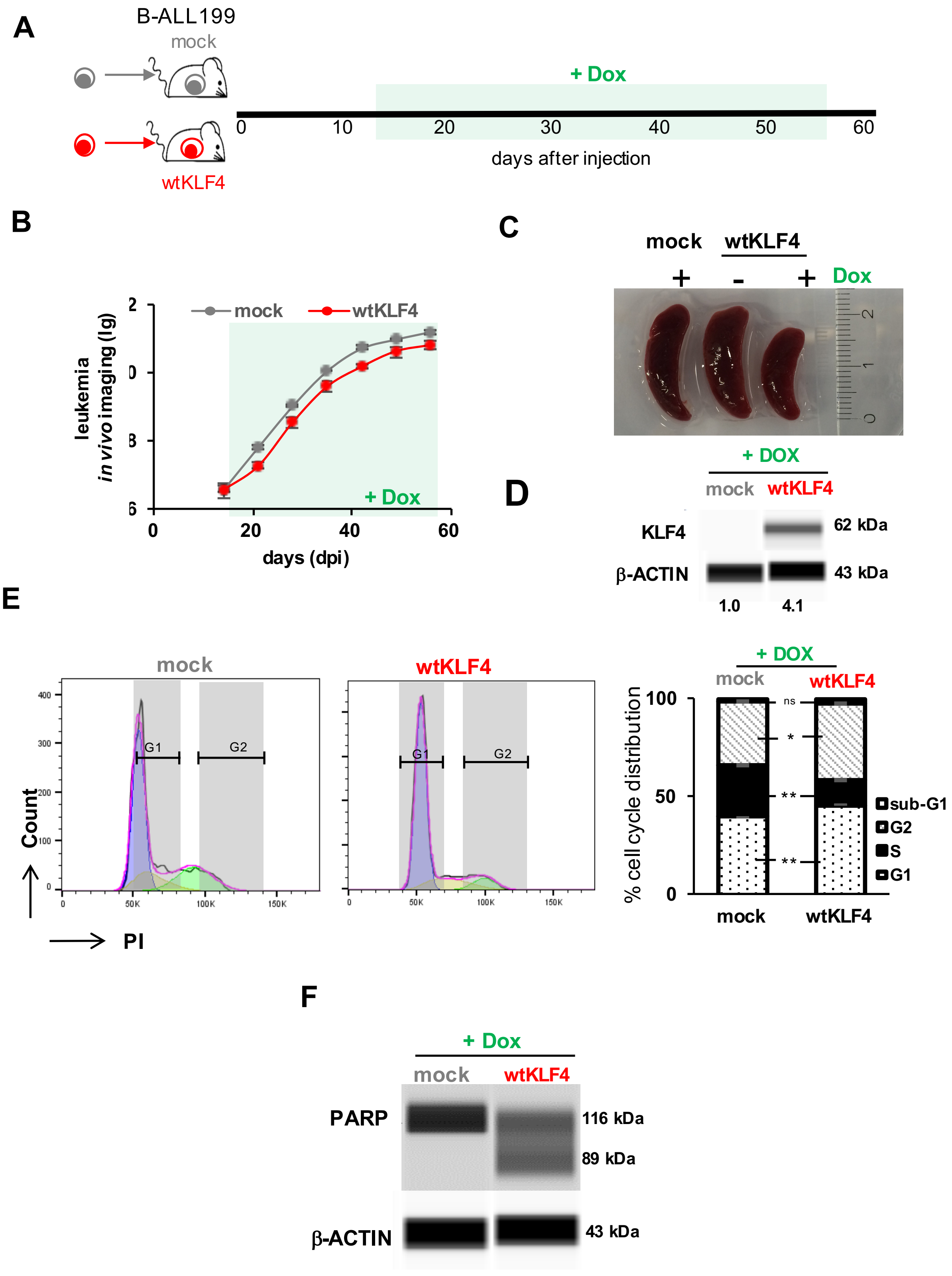

A
B-ALL199
mock
+ Dox
0 10 20 30 40 50 60
days after injection
wtKLF4
B
leukemia
in vivo imaging (lg)
mock
wtKLF4
+ Dox
0 20 40 60
days (dpi)
C
mock
wtKLF4
+ - +
Dox
D
+ DOX
mock wtKLF4
KLF4
62 kDa
β-ACTIN
43 kDa
1.0 4.1
E
mock
wtKLF4
Count
G1 G2
PI
+ DOX
mock wtKLF4
% cell cycle distribution
100
50
0
ns
*
**
**
sub-G1
G2
S
G1
mock wtKLF4
F
+ Dox
mock wtKLF4
PARP
116 kDa
89 kDa
β-ACTIN
43 kDa

**Figure S4 Re-expressing KLF4 inhibits tumor growth through cell cycle arrest and apoptosis in B-ALL PDX cells *in vivo***

Experiments were performed and depicted identically as in printed Figure 3, except that ALL-199 was used instead of ALL-265. ALL-199 cells were injected into 6 NSG mice each. After homing was completed and tumors were established, DOX was added to the drinking water on day 14 and leukemia growth monitored by bioluminescence *in vivo* imaging. On day 56 after cell injection, mice were sacrificed, spleens harvested and the successfully DOX-induced, mCherry positive mock or wtKLF4 expressing population enriched by flow cytometry for further analysis.

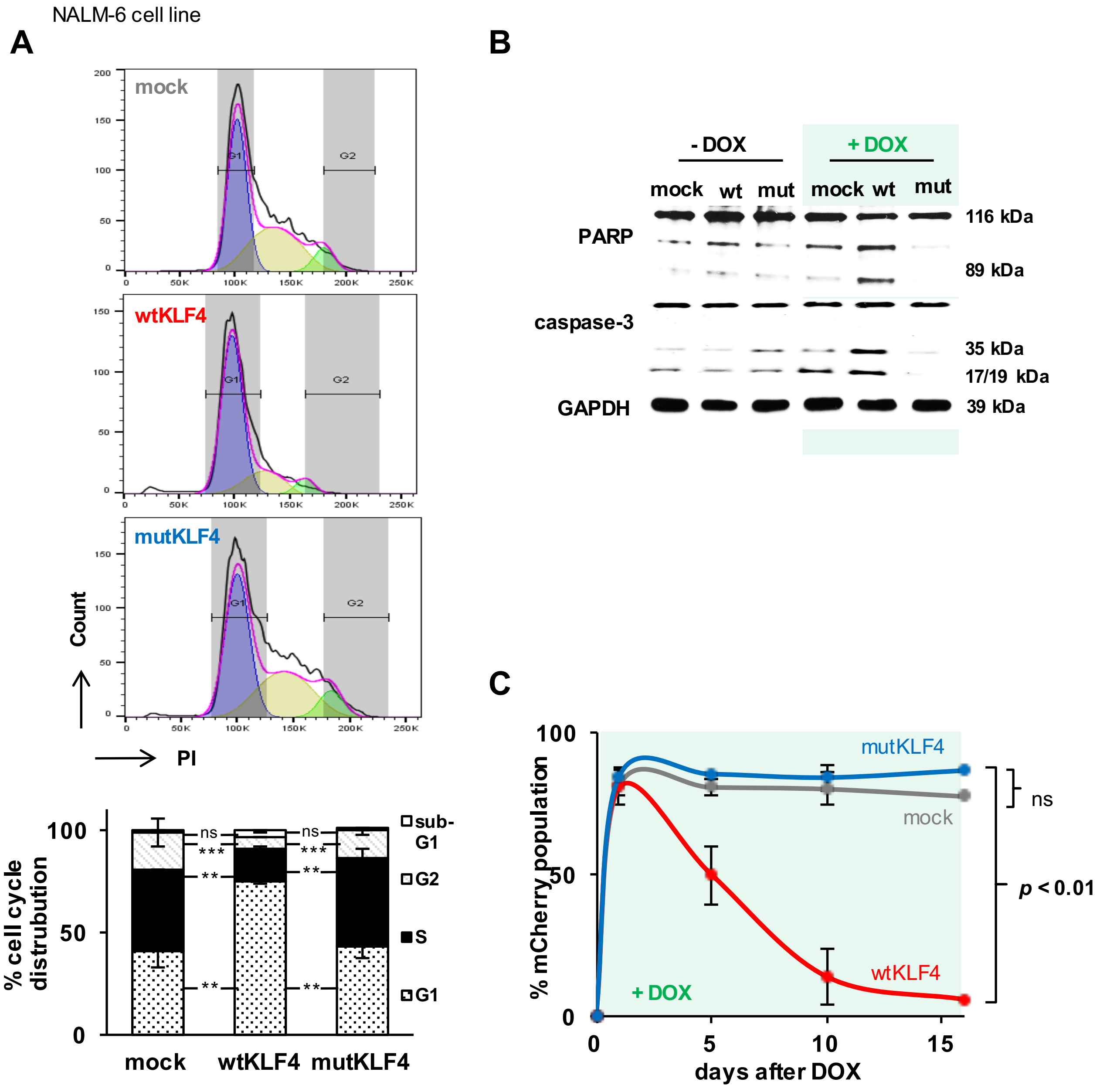


**Figure S5** **Re-expressing KLF4 inhibits tumor growth through cell cycle arrest and apoptosis in NALM-6 cells *in vitro***

$10^6$ NALM-6 cells were transduced with the constructs indicated (Figure S1C) and cultured in the presence of 0.5µg/ml DOX.

A. After 2 days, cells were harvested and analyzed and depicted as in Figures 3E and S4E. ** p<0.01, *** p<0.005 as determined by two-tailed unpaired t test.

B. Western Blot was performed as in Figures 3F and S4F.

C. mCherry expression indicating expression of mock, wtKLF4 or mut KLF4 was monitored by flow cytometry at the indicated time points; mean ± SD of 3 independent experiments is shown. ns: not significant, *** p<0.005 as determined by two-tailed unpaired t test.

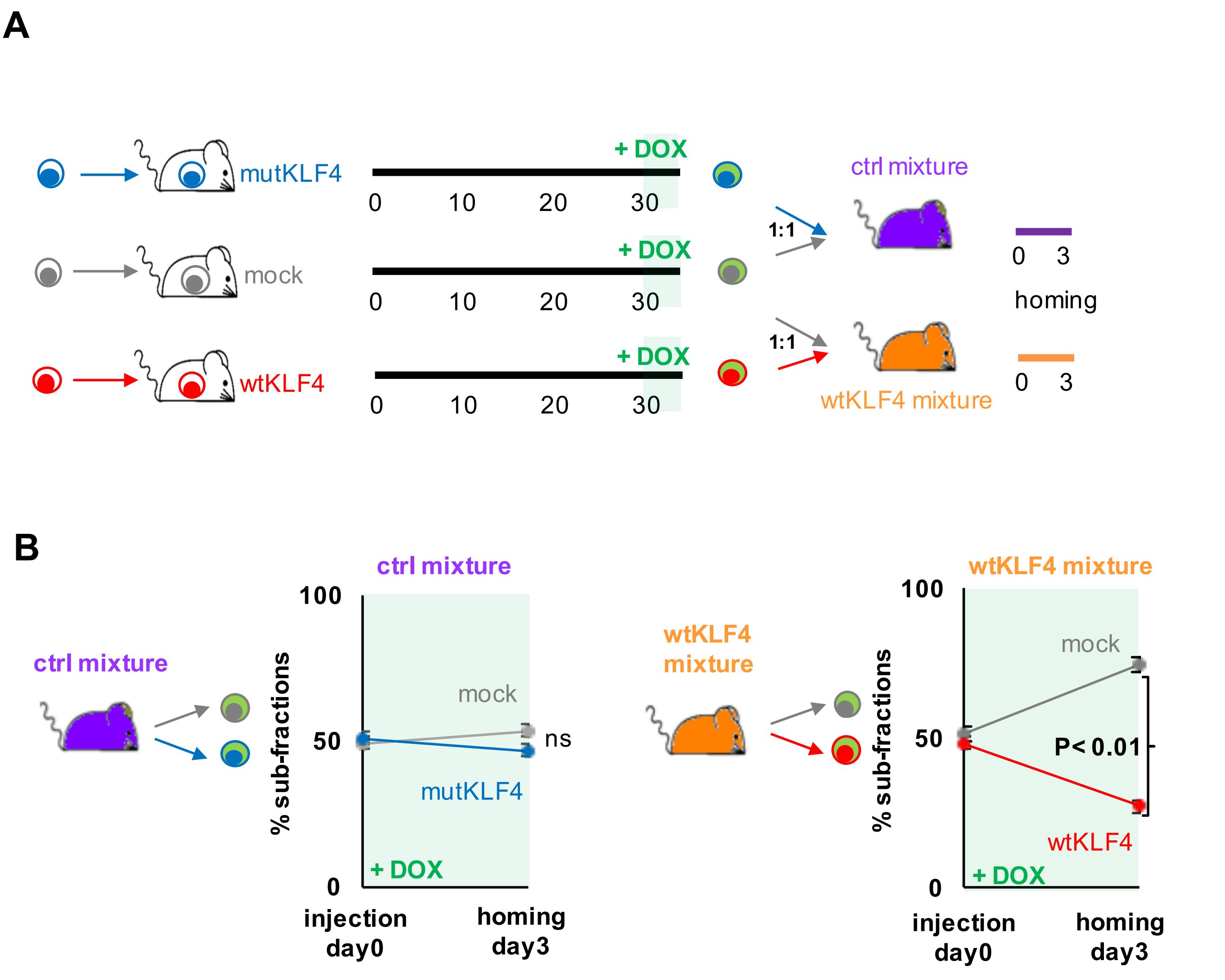


**Figure S6 wtKLF4 inhibits homing of PDX ALL cells to the murine bone marrow**

A. Experimental procedure: ALL-265 cells transduced with either mock, mutKLF4 or wtKLF4 constructs were separately injected into NSG mice and DOX administered during the last 48 hours before mice were sacrificed. Successfully induced, mCherry positive cells were enriched and mixed into either control or KLF4 mixture, identically as in printed Figures 4 and 5. $10^6$ cells of each mixture were re-injected into 3 NSG recipients each. To evaluate homing capacities, bone marrow of secondary recipients was harvested 3 days after injection and analyzed by flow cytometry.

B. Subfraction analysis was performed and is depicted as mean ± SEM of 3 mice per group. ns: not significant; * $p < 0.01$ as determined by two-tailed unpaired t test.

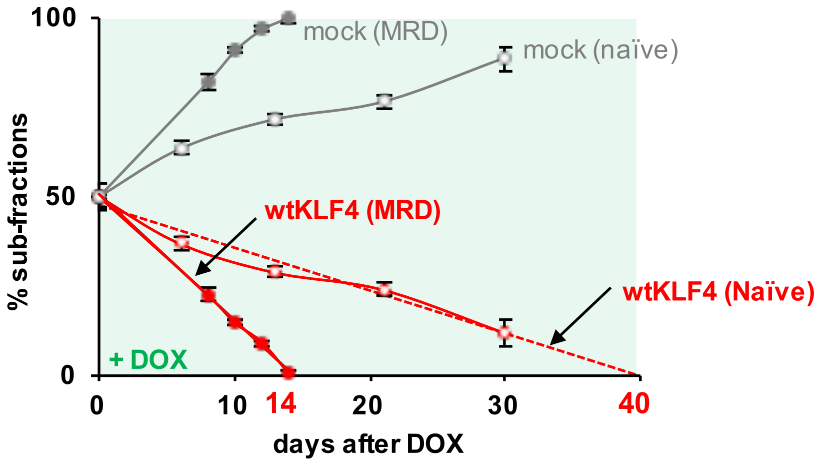


**Figure S7 Treatment-surviving cells are especially sensitive towards re-expressing KLF4**

Graph compares data from sub-fractions of wtKLF4 mixture from Figure 4B (untreated cells) with Figure 4D (treated cells). Red dashed line represents calculated trend of untreated cells.

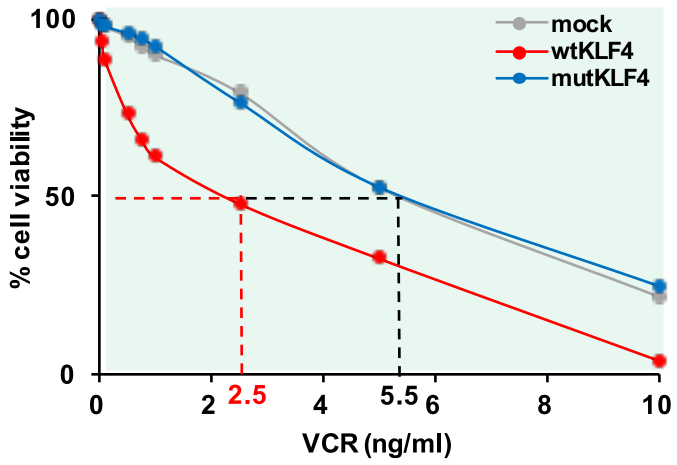


**Figure S8** **Re-expressing KLF4 sensitizes NALM-6 cells towards Vincristine treatment in vitro**

$10^6$ NALM-6 cells transduced with the indicated constructs were cultured in the presence of 0.5µg/ml DOX and treated with increasing concentration of Vincristine (VCR) for 48h. Cell viability was analyzed by flow cytometry; all groups were normalized to non-VCR treated cells; mean ± SD of 3 independent experiments is shown.

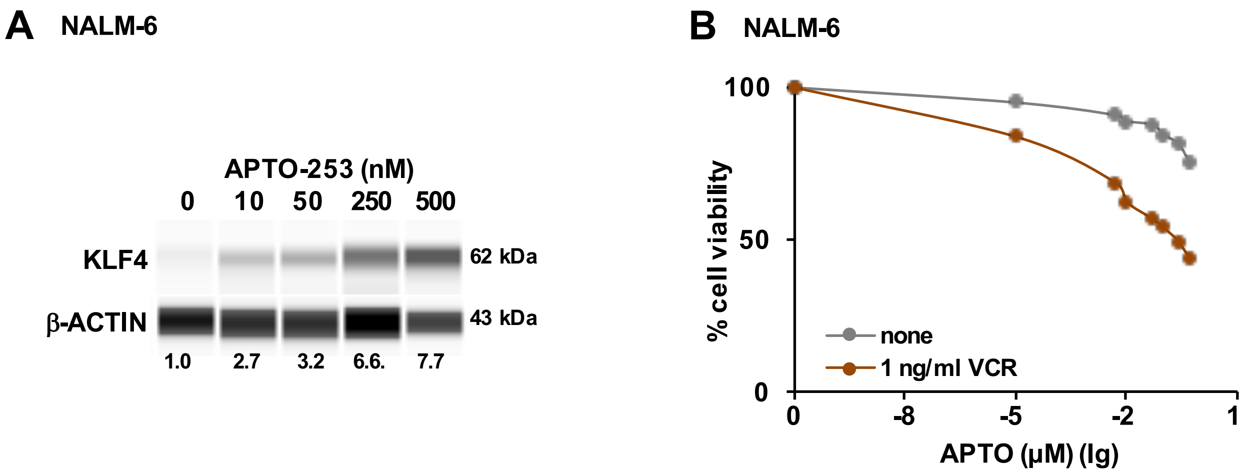


**Figure S9 APTO-253 upregulates KLF4 and enhances sensitivity to Vincristine**

A. NALM-6 cells were incubated with the indicated concentrations of APTO-253 for 3 days and KLF4 protein expression analyzed by capillary immunoassay; beta-actin was used as loading control; one representative immunoblot of 3 independent experiments is shown.

B. NALM-6 cells were incubated with the indicated concentrations of APTO-253 in the presence or absence of 1ng/ml VCR. After 48h, cell viability was analyzed by flow cytometry; mean ± SD of 3 independent experiments is shown.

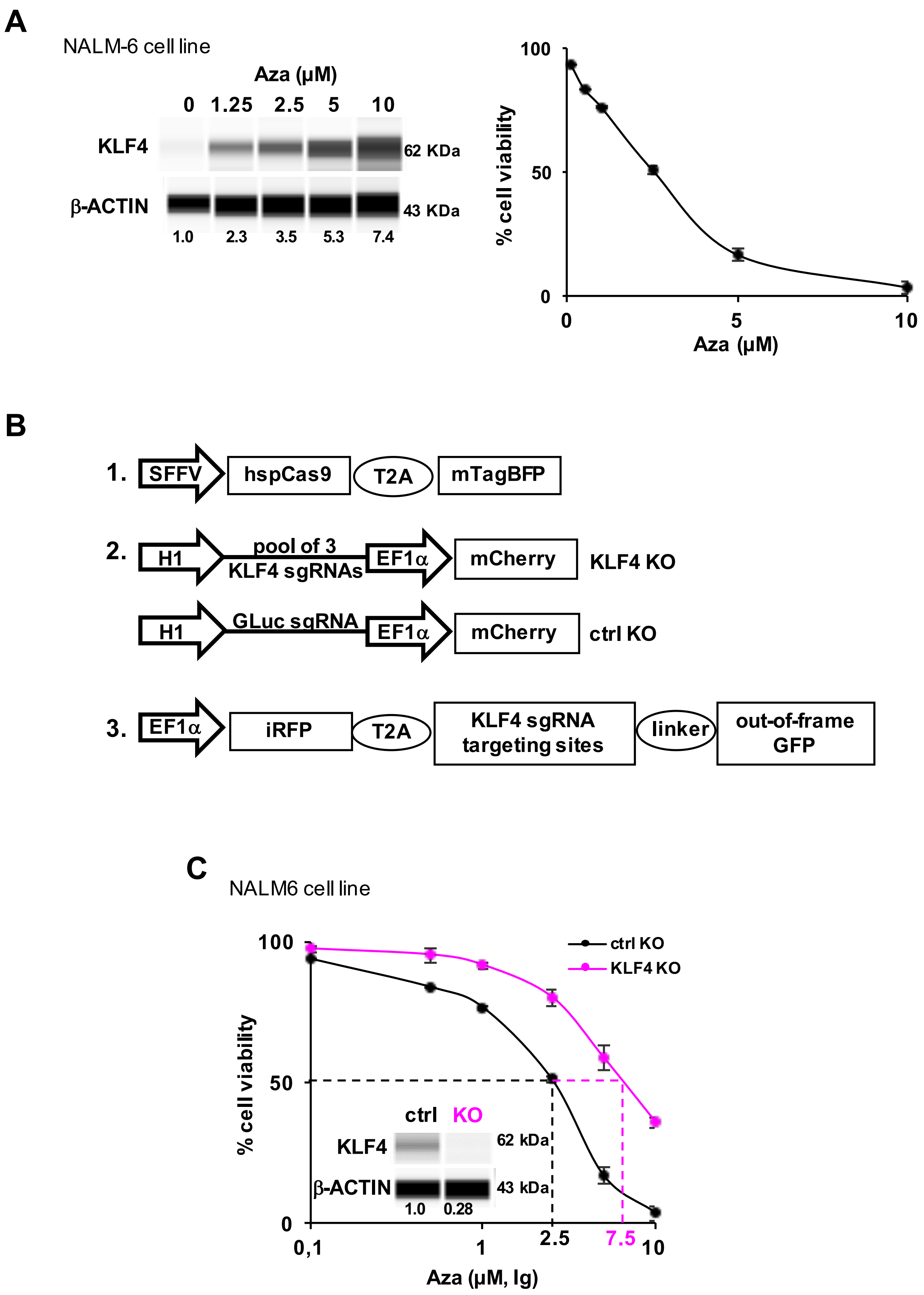
A
NALM-6 cell line
Aza (µM)
0 1.25 2.5 5 10
KLF4
62 KDa
β-ACTIN
43 KDa
1.0 2.3 3.5 5.3 7.4
% cell viability
100
50
0
0 5 10
Aza (µM)
B
1.
SFFV
hspCas9
T2A
mTagBFP
2.
H1
pool of 3 KLF4 sgRNAs
EF1α
mCherry
KLF4 KO
H1
GLuc sgRNA
EF1α
mCherry
ctrl KO
3.
EF1α
iRFP
T2A
KLF4 sgRNA targeting sites
linker
out-of-frame GFP
C
NALM6 cell line
100
50
0
% cell viability
ctrl KO
KLF4 KO
ctrl KO
KLF4
62 kDa
β-ACTIN
43 kDa
1.0 0.28
0,1 1 2.5 7.5 10
Aza (µM, lg)

**Figure S10 Azacitidine-induced cell death depends on upregulating KLF4**

A. NALM-6 cells were treated with different concentrations of Azacitidine (Aza) for 48h *in vitro.*

1. Left panel: KLF4 protein expression analyzed by capillary immunoassay, beta-actin served as loading control. One representative immunoblot out of 2 independent experiments is shown.
2. Right panel: Viability was analysed by flow cytometry.

B. Schematic diagram of expression constructs for knockout:

1. A human codon-optimized S. pyogenes (hsp)Cas9 was constitutively expressed from the SFFV promoter together with the fluorochrome mTaqBFP, linked via a T2A sequence.
2. The single-guide (sg)RNA was expressed under the control of the H1 Pol III promoter. 3 different KLF4 sgRNAs were cloned individually into the constructs before vectors were pooled. GLuc sgRNA was used as a control. mCherry expression from the EF1a promoter allowed enriching transgenic cells.
3. A reporter construct was used for enriching KLF4 knockout cells; in brief, EF1a drives constitutive expression of iRFP720 as well as out-of-frame GFP 5′ of a PAM-containing Cas9/sgRNA targeting sequences. sgRNA guided gene editing in the targeting sequence by Cas9 shifts GFP into frame so that GFP expressing cells are strongly enriched for KLF4 gene knockout. All the constructs were cloned into the pCDH backbone.

C $10^6$ KLF4 KO or ctrl NALM-6 cells were treated with the indicated concentrations of Aza for 2 days and cell viability was analyzed by flow cytometry. Cell viability was normalized to untreated cells. Dashed lines indicates IC50 of ctrl KO (black) and KLF4 KO (pink) cells. Mean ± SD of 3 independent experiments is shown. Inner panel: KLF4 protein level of NALM-6 cells were detected by capillary immunoassay.
